# Methionine Restriction Reduces Lung Cancer Progression and Increases Chemotherapy Response

**DOI:** 10.1101/2024.06.25.599795

**Authors:** Kassandra J. Naughton, Xiulong Song, Avery R. Childress, Erika M. Skaggs, Aria L. Byrd, Christian M. Gosser, Dave-Preston Esoe, Tanner J. DuCote, Daniel R. Plaugher, Alexsandr Lukyanchuk, Ryan A. Goettl, Jinpeng Liu, Christine F. Brainson

## Abstract

Targeting tumor metabolism through dietary interventions is an area of growing interest, and may help to improve the significant mortality of aggressive cancers, including non-small cell lung cancer (NSCLC). Here we show that the restriction of methionine in the aggressive KRAS*/Lkb1-*mutant NSCLC autochthonous mouse model drives decreased tumor progression and increased carboplatin treatment efficacy. Importantly, methionine restriction during early stages of tumorigenesis prevents the lineage switching known to occur in the model, and alters the tumor immune microenvironment (TIME) to have fewer tumor-infiltrating neutrophils. Mechanistically, mutations in *LKB1* are linked to anti-oxidant production through changes to cystathionine-β-synthase (CBS) expression. Human cell lines with rescued *LKB1* show increased CBS levels and resistance to carboplatin, which can be partially rescued by methionine restriction. Furthermore, LKB1 rescued cells, but not mutant cells, show less G2- M arrest and apoptosis in high methionine conditions. Knock-down of CBS sensitized both LKB1 mutant and non-mutated lines to carboplatin, again rescuing the carboplatin resistance of the LKB1 rescued lines. Given that immunotherapy is commonly combined with chemotherapy for NSCLC, we next wanted to understand if T cells are impaired by MR. Therefore, we examined the ability of T cells from MR and control tumor bearing mice to proliferate in culture and found that T cells from MR treated mice had no defects in proliferation, even though we continued the MR conditions *ex vivo*. We also identified that CBS is most highly correlated with smoking, adenocarcinomas with alveolar and bronchiolar features, and adenosquamous cell carcinomas, implicating its roles in oxidative stress response and lineage fate in human tumors. Taken together, we have shown the importance of MR as a dietary intervention to slow tumor growth and improve treatment outcomes for NSCLC.

## INTRODUCTION

Methionine restriction (MR) diets have gained attention due to reports that cancers are specifically sensitive to reduced dietary methionine, and that MR potentiates both chemotherapy and radiation responses [1, 2]. As an essential amino acid, methionine is critical for protein production, and is the key player in the methionine cycle that produces methyl groups for DNA, histones, proteins, RNA, and lipid methylation [3]. In this cycle, methionine is converted into S-adenosyl-methionine (SAM) via addition of ATP by the enzyme methionine adenosyltransferase (MAT2A). SAM-dependent methyltransferases can then cleave the sulfur-bound methyl group and subsequently methylate substrates, leaving behind S-adenosyl-homocysteine (SAH). SAH functions as a precursor to homocysteine, and acts as an inhibitor of SAM-dependent methyltransferases [3, 4]. The conversion of SAH into homocysteine leads to a branch point in the cycle. Homocysteine can either be recycled back to methionine via donors from the folate cycle, or shunted towards cytathionine by the rate-limiting activity of cystathionine-β-synthase (CBS) [3, 5]. This begins the transsulfuration pathway, that can ultimately to produce glutathione, an important antioxidant in reactive oxygen species (ROS) mitigation [3]. Once this conversion from homocysteine to cystathionine occurs, it is irreversible, thus limiting methionine recycling and in MR conditions, and possibly reducing pools of both methionine and SAM for methylation.

For maintenance of epigenetic states, one key SAM-dependent methyltransferases is Enhancer of Zeste Homolog 2 (EZH2) within the Polycomb Repressive Complex 2 (PRC2). PRC2 uses SAM to tri-methylate Histone 3 at lysine 27 (H3K27me3), and this chromatin mark leads to silenced gene expression [6]. EZH2 is highly expressed in many tumor types, and is known to silence tumor suppressors and genes involved in antigen presentation [6, 7]. In non-small cell lung cancers (NSCLCs) H3K27me3 levels can vary, with lung adenocarcinomas (ADCs) overall showing higher levels of H3K27me3 than lung squamous cell carcinomas (SCCs) [8, 9]. Intriguingly, cystathionine levels are also higher in SCCs, suggesting that an increase in the transsulfuration pathway could limit SAM levels and ultimately lead to the decreased H3K27me3 observed in some SCCs [8–10].

In addition to epigenetic differences, genetic heterogeneity of NSCLCs contributes to tumor phenotypes and drug resistance. For example, while *KRAS*-driven tumors are particularly aggressive and respond poorly to immunotherapy, those with additional serine-threonine kinase 11 (*STK11* aka *LKB1)* mutation have even worse outcomes [11–13]. The major function of LKB1 is to phosphorylate and activate AMPK in response to a high AMP/ATP ratio, indicating low levels of energy (in the form of ATP) [14]. When *LKB1* is mutated, this inhibits the ability of cells to sense changing AMP/ATP ratios and respond to metabolic stressors efficiently. It also means that pathways that utilize ATP, including SAM production, are not properly regulated. It has been observed that in both murine and human NSCLCs, levels of SAM are lower in LKB1-deficient tumors compared to LKB1 intact tumors [10, 15]. We previously demonstrated that, in an aggressive model of KRAS/*Lkb1* NSCLC, tumor progression from ADC to SCC is modulated in part through loss of H3K27me3 gene silencing at key SCC lineage genes including *Sox2* and *Trp63*. This squamous transdifferentiation is only observed when LKB1 is mutated, again suggesting that inability of LKB1 mutant tumors to maintain proper SAM levels could drive the loss of PRC2 activity and the epigenetic shift [8].

Several studies in colorectal cancer, sarcoma and melanoma have demonstrated that methionine restriction potentiates therapies including radiation and chemotherapy [1, 16]. However, the models used were all LKB1 proficient, and it remains unclear if KRAS/*Lkb1* NSCLC would be similarly impaired by MR. Here, we use a combination of autochthonous mouse models, human lung cancer cell lines and patient samples to learn if MR is an effective treatment strategy for KRAS/*Lkb1* NSCLCs, and to learn how MR influences methionine metabolism and histone methylation in LKB1-deficient settings.

## RESULTS

### Methionine restriction to prevent tumor progression

We first sought to determine if restricting dietary methionine can reduce the proliferation of LKB1-deficient lung cancer. To achieve this, we used the autochthonous genetically engineered mouse model harboring oncogenic Cre-inducible KRAS^G12D^ and *Lkb1* floxed alleles (*Kras^mut^/Lkb1^mut^*). One week post intranasal instillation of adenoviral-Cre, mice were split into two equal groups of control (0.86% w/w) and methionine restricted (0.12% w/w) (MR) diets for 9 weeks (**Figure 1A**). All mice had weight reduction after adeno-Cre, and while the control chow group regained this weight, the MR group remained at stable weights at 91% of initial (**Supp.** Figure 1A). To confirm the intended decrease in methionine levels with the MR diet, the serum of both control and MR mice was analyzed for both methionine and SAM concentrations (**Figures 1B+C**). Both methionine and SAM serum concentrations were significantly decreased in MR mice compared to the control mice. These data suggest that the MR diet is not only sustainable and safe for mice over an extended period but also indicates the efficacy of the diet to reduce serum methionine and SAM levels.

**Figure 1:**
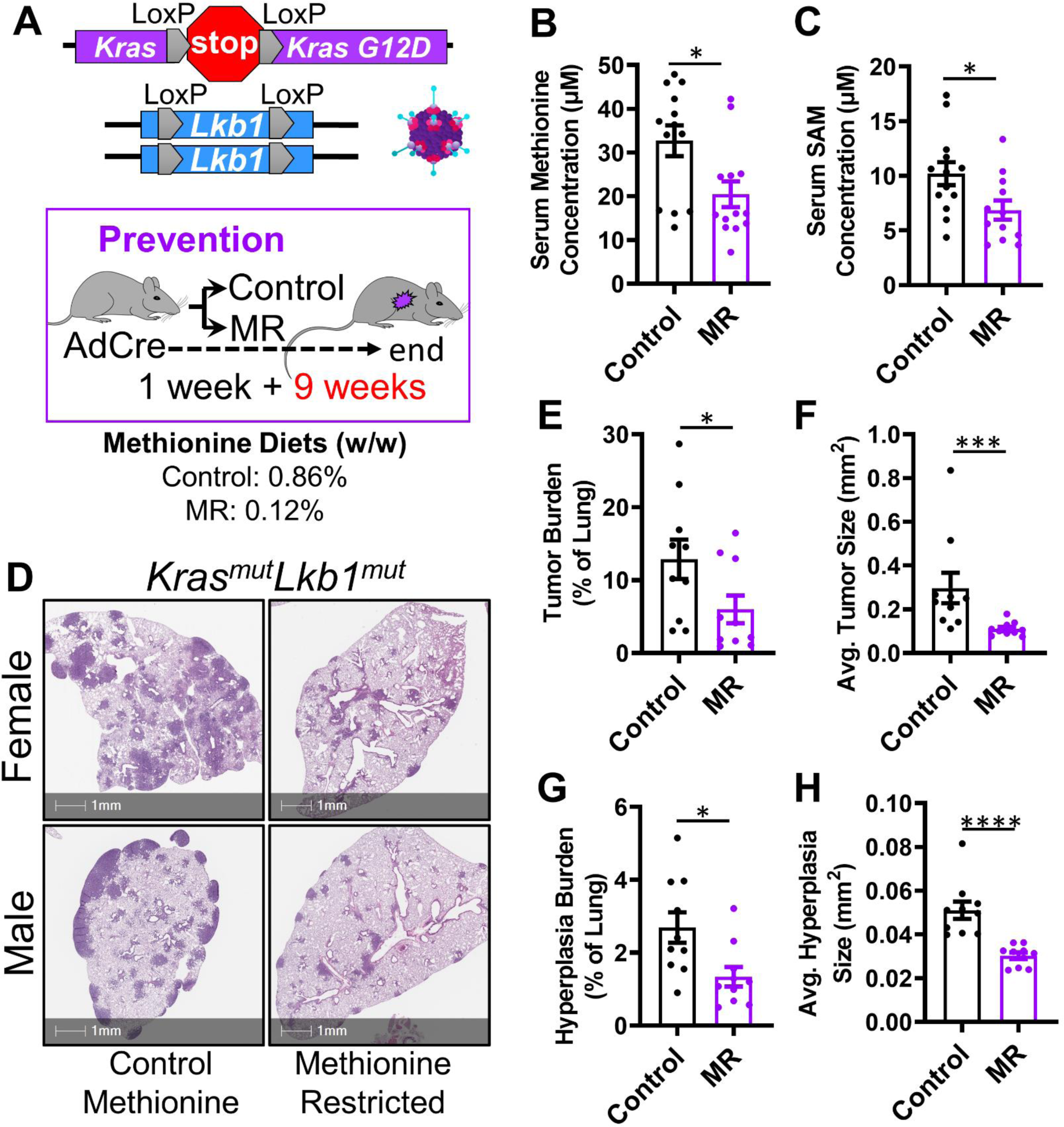
Methionine Restriction Reduces Tumor Growth in the KRAS/*Lkb1* Lung Cancer Model **A)** Schematic of alleles and dietary intervention for tumor progression prevention. *KrasG12D* activation and *Lkb1* deletion occur in a single dose of Adeno-Cre Virus administered intranasally to mice. One week post administration, mice were separated into control methionine (0.86% w/w) and methionine restricted (0.12% w/w) dietary groups for 9 weeks. **B)** Methionine concentration in serum of mice placed on control and methionine restricted diets plotted as mean +/- SEM. n=13 for control and methionine restricted groups, which includes 10 tumor-bearing mice and 3 non-tumor bearing mice for each group, * indicates p=0.0102 by two-tailed Mann-Whitney U test. **C)** SAM concentration in serum of mice placed on control and methionine restricted diets plotted as mean +/- SEM. n=13 for control, which includes 10 mice with tumors and 3 control mice without tumors, and n=12 for methionine restricted group which includes 9 mice with tumors and 3 control mice without tumors. One tumor-bearing mouse was removed from the MR group after performing the ROUT outlier identification test, * indicates p=0.0257 by two-tailed Mann-Whitney U test. **D)** Representative Hematoxylin and Eosin (H+E) staining images of male and female lungs of *KrasG12D*-activated and *Lkb1*-null mice on control and methionine restricted diets, scale bars=1mm. **E)** Tumor burden of control and methionine restricted mice calculated based on the percent of total lung area that was identified as tumor, plotted as mean +/- SEM, n=10 for control and methionine restricted groups, * indicates p=0.0355 by Two-tailed Mann-Whitney U test. **F)** Average tumor size in mm^2^ of all tumors identified in control and methionine restricted groups plotted as mean +/- SEM, n=10 mice per group, each data point representing average tumor area of each mouse, *** indicates p=0.0002 by two-tailed Mann-Whitney U test. **G)** Hyperplasia burden of control and methionine restricted mice calculated based on the percent of total lung that was classified as hyperplasia, plotted as mean +/- SEM, n=10 for control and methionine restricted groups, * indicates p=0.0147 by two-tailed Mann-Whitney U test. **H)** Average hyperplasia size in mm^2^ of all hyperplasia identified in control and methionine restricted groups plotted as mean +/- SEM, n=10 mice per group, each data point representing average tumor area of each mouse, **** indicates p<0.0001 by two-tailed Mann-Whitney U test.

After the 9 weeks on MR or normal chow and 10 weeks after adeno-Cre, all five lobes of the lungs of the mice were inflated with formalin, embedded, and sectioned for histological analysis (**Figure 1D**). The lungs of a group of *Kras*^WT^*/Lkb1*^WT^ on the same control and MR diets were also imaged and showed no tumors and no signs of inflammation or fibrosis in the MR group (**Supp.** Figure 1B). In the tumor-bearing group, there was no significant difference in the number of tumors between the MR and normal chow cohorts (**Supp.** Figure 1C). This was expected given that the adeno-Cre virus was given to initiate the tumors prior to the start of the MR diet. Despite no significant difference in the number of tumors that formed, MR mice had significantly lower tumor burden, calculated as the percent of total lung identified as tumor (**Figure 1E**), and significantly lower average tumor size (**Figure 1F**), as compared to the control methionine mice. This suggests that while MR does not decrease tumor initiation (total number of tumors), it instead slows the progression of those tumors (tumor burden and size). Hyperplasia can also be an indicator of potential tumor development, and we found that both hyperplasia burden (**Figure 1G**) and hyperplasia size (**Figure 1H**) were significantly lower in the MR mice compared to mice on normal chow. During the experiment, we also noticed that the female mice tended to have higher overall tumor burden. When separated by sex, the data show that methionine restriction works for both sexes, but that females have higher tumor burden overall, and therefore the most striking tumor suppression was observed in the MR males (**Supp.** Figure 1D). Together, these data suggest an important role for MR in slowing tumor progression and development in the aggressive *Kras^mut^/Lkb1^mut^* lung cancer model.

To further explore the role of methionine during oncogenic transformation, we utilized the human BEAS-2B lung cell model that already has a dominant negative p53 (TP53-R175H) and can be transformed by oncogenic RAS [17]. BEAS-2B cells sensitized in high (574.9 µM), regular (115.3 µM) or low (57.9 µM) methionine media were transduced with a Kras-G12V virus and selected for viral infection. We chose the low methionine concentration based on the observation that *in vivo* MR diets can decrease the serum levels of methionine by only half [1] (**Figure 1B**). These cells were then transferred into high, regular, and low methionine media in soft agar to produce tumoroids. Cells that were transduced with virus in high methionine had no changes in tumoroid number when grown in the other concentrations of methionine in the soft agar (**Supplementary** Figure 1E). However, cells that were transformed in either regular or low methionine media and then grown in soft agar in high methionine had significantly more tumoroids form than those grown in soft agar in either regular or low methionine media. Conversely, cells kept in low methionine media during the growth phase were unable to grow as many colonies in the soft agar. Together, these data suggest that when transformation occurs in a methionine-depleted environment, high methionine can increase cell proliferation or survival.

### Methionine restriction changes tumor phenotype and immune microenvironment

Several laboratories, including our own, have described the progression of the *Kras*^mut^/*Lkb1*^mut^ mouse model from primarily ADCs to SCCs [8, 18, 19]. From a stem cell perspective, an intermediary between an alveolar type 2 driven ADC and an SCC would be a bronchiolar cell phenotype [20]. Therefore, to determine if MR significantly alters the lineage phenotypes, all histologically identified tumors were further classified into either alveolar adenocarcinoma (A-ADC), bronchiolar adenocarcinoma (B-ADC), or SCC tumor based on cellular morphology in H+E-stained slides (**Figure 2A**). We then compared control methionine to MR mice tumor phenotypes and found that although A-ADC tumor burden and tumor number were not significantly decreased by MR (**Figure 2B, Supp.** Figure 2A), there was a significant reduction in the average size of A-ADC tumors in MR (**Figure 2C**). In addition, B-ADC tumor number, tumor burden, and tumor size were all significantly decreased in the MR mice (**Figure 2D+E, Supp.** Figure 2B). Finally, we examined SCC tumors, which should arise in 40% of the mice. While 4 of 10 mice in the control chow group had SCC tumors, none of the mice in the MR group had this tumor phenotype (**Supp.** Figure 2C). These results suggest that MR may be reducing the progression of tumors from A-ADCs to B-ADC and SCC tumors.

**Figure 2:**
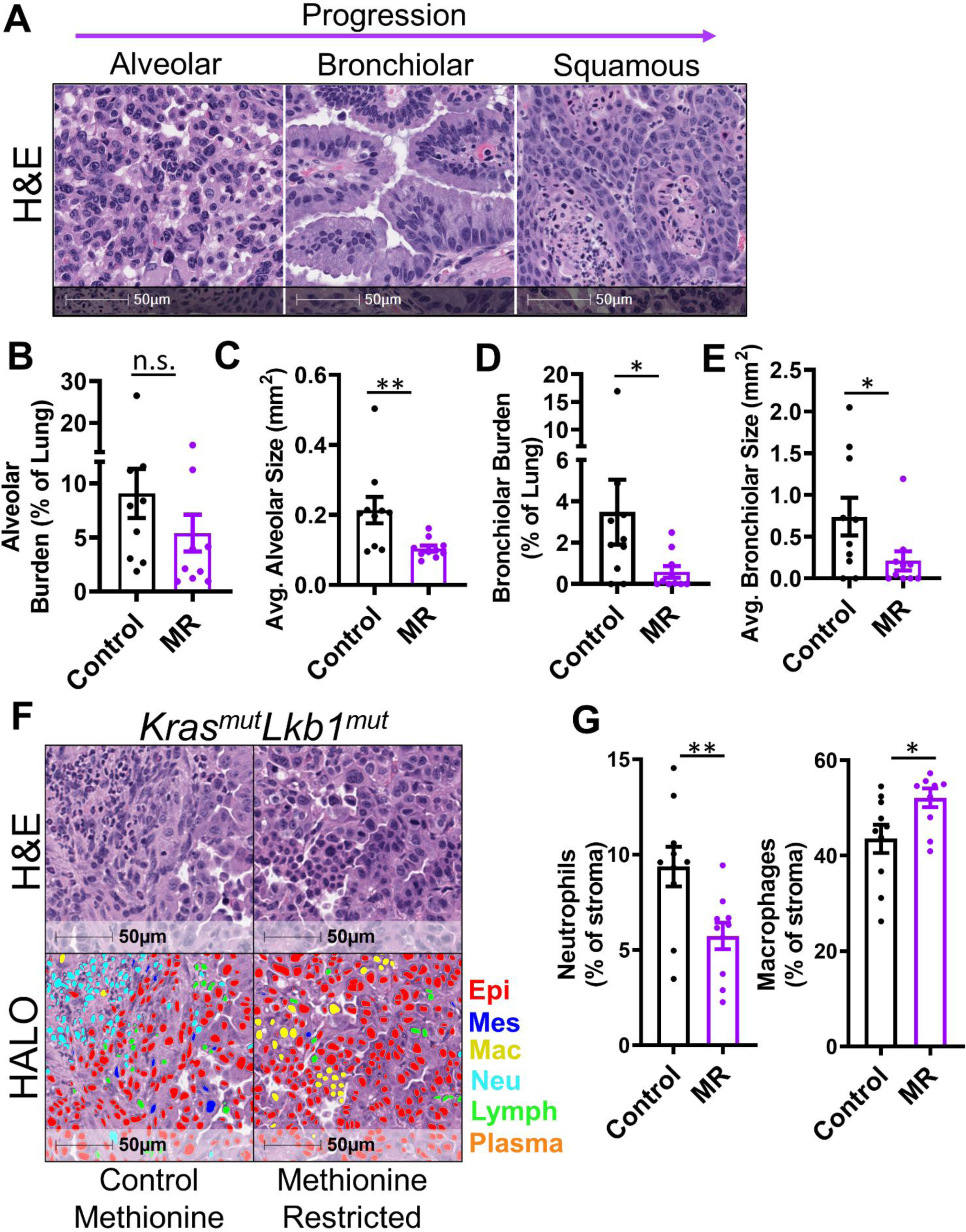
Methionine Restriction Reduces Tumor Progression and Alters Tumor Immune Cells **A)** Representative H+E staining images of alveolar, bronchiolar, and squamous tumors from the *Kras^G12D^/Lkb1^null^* mice, scale bars=50µm. Purple progression arrow indicates projected linage progression of *Kras/Lkb1*-mutated murine tumors. **B)** Alveolar tumor burden in control and methionine restricted mice, calculated based on the percent of total lung area that was identified as alveolar tumor, plotted as mean +/- SEM, n=10 mice per group, n.s.=not significant by two-tailed Mann-Whitney U test. **C)** Average alveolar size in mm^2^ of all tumors identified as alveolar for control and methionine restricted mice, plotted as mean +/- SEM, n=10 mice per group, each data point representing average tumor area of each mouse, ** indicates p=0.0029 by two-tailed Mann-Whitney U test. **D)** Bronchiolar tumor burden in control and methionine restricted mice, calculated based on the percent of total lung area that was identified as bronchiolar tumor, plotted as mean +/- SEM, n=10 mice per group, * indicates p=0.0311 by Two-tailed Mann-Whitney U test. **E)** Average bronchiolar tumor size in mm^2^ of all tumors identified as alveolar for control and methionine restricted mice, plotted as mean +/- SEM, n=10 mice per group, each data point representing average tumor area of each mouse, * indicates p=0.0382 by Two-tailed Mann-Whitney U test. **F)** Representative H+E staining and HALO® nuclear phenotyper images of control and methionine restricted mouse tumors. Nuclear phenotyper identifies immune cells in the tumor immune microenvironment. Epithelial (Epi) (or tumor cells) are marked in red, mesenchymal (Mes) cells are dark blue, macrophages (Mac) are in yellow, neutrophils (Neu) are in light blue, lymphocytes (Lymph) are in green, and plasma cells are in orange. Scale bars=50µm. **G)** Percent of neutrophils and macrophages of all non-epithelial cells in the immune microenvironment of control and methionine restricted mice, plotted as mean +/- SEM, n=10 mice per group, * indicates p=0.0258, ** indicates p=0.0094 by two-tailed t-test.

We and others have previously demonstrated that SCC transdifferentiation in this model is accompanied by a large influx of tumor-associated neutrophils [8, 19, 21]. After annotating tumors as the different lineage phenotypes, a trained and verified macro on the HALO® nuclear phenotyper was applied to the H+E slides [22].

This nuclear phenotyper identified different immune cells in the tumor immune microenvironment (TIME), including epithelial cells (Epi, red), mesenchymal cells (Mes, dark blue), macrophages (Mac, yellow), neutrophils (Neu, light blue), lymphocytes/monocytes (Lymph, green), and plasma cells (Plasma, orange). Consistent with previously described immune infiltrating cells in tumors of different phenotypes [10, 22], A-ADCs had a large proportion of macrophages, while SCCs had larger proportions of neutrophils (**Supp.** Figure 2B). A representative image of a tumor from a control chow mouse and one from an MR mouse show the differences in the tumor immune microenvironment adjacent to the H+E stain (**Figure 2F**). In MR treated mice, the number of infiltrating macrophages was significantly higher, while the number of infiltrating neutrophils was significantly in all tumors when compared to tumors from normal chow mice (**Figure 2G**).

### LKB1 expression alters methionine metabolism in human NSCLC cells

The methionine and transsulfration cycles have numerous metabolites, enzymes and transporters that regulate its activity (**Figure 3A**). To dissect the roles of LKB1 in these cycles, we began by examining the levels of CBS in LKB1 wild-type (LKB1-WT) and LKB1-mutant NSCLC cell lines grown in differing methionine conditions. Western blot of two *KRAS*^mut^/*LKB1*^mut^ NSCLC cell lines (A549 and H2030) and one *KRAS*^mut^/*LKB1*^WT^ NSCLC cell line (H2009) grown in high, regular, and low methionine media concentrations showed much higher levels of CBS in the H2009 LKB1-WT cell line than in the two mutant lines (**Figure 3B**). Relative expression of *CBS* mRNA confirmed higher levels of *CBS* in the LKB1-WT H2009 cells (**Figure 3C**). Low methionine could reduce CBS protein levels. However, no significant differences were seen in different concentrations of methionine, which matches reports that methionine stabilizes CBS at the protein level [23]. Next, to confirm differences seen between *LKB1*-proficient and *LKB1*-mutant cells, we rescued LKB1 in A549 cells (**Figure 3D+E**). Consistent with the parental blots, CBS expression was higher in LKB1-rescue A549, regardless of methionine levels (**Figure 3D, Supp.** Figure 3A). CBS levels were the lowest in LKB1-mutant cells grown in low methionine, and these cells also had a 2-fold reduction in H3K27me3 (**Figure 3D, Supp.** Figure 3A). These data suggest MR drives a stronger loss of SAM pools when LKB1 is mutated, likely due to lower availability of ATP to form new SAM molecules.

**Figure 3:**
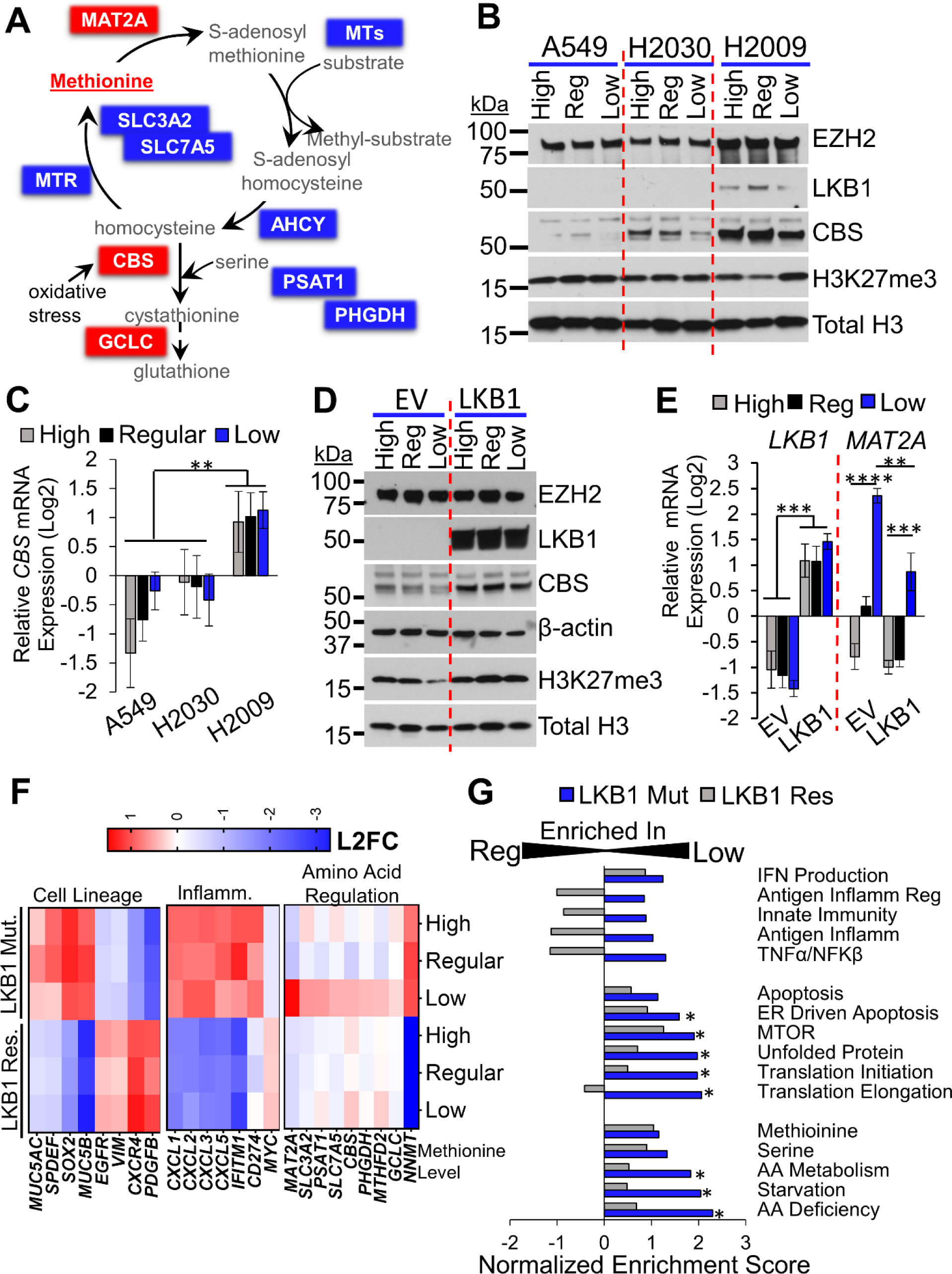
LKB1 Changes Methionine Restriction Response and PRC2 Activity **A)** Schematic of the methionine cycle. Methionine is either obtained from exogenous sources through transporters (SLC3A2 and SLC7A5), salvaged in the methionine salvage pathway, or recycled in the methionine cycle. MAT2A first converts methionine into S-adenosyl methionine (SAM), the universal methyl donor. Methyl groups are cleaved from SAM via various methyl transferases (MT) and converted into S-adenosyl homocysteine (SAH). MTs deposit methyl groups onto DNA, histones, etc. SAH is further converted into homocysteine via adenosylhomocysteine hydrolase (AHCY). Homocysteine is either recycled back into methionine via methionine synthase and donors from the folate cycle or used in the transsulfuration pathway to produce glutathione through conversion into cystathionine via cystathionine-β-synthase (CBS). Serine biosynthesis plays an important role in providing serine for use in the transsulfuration pathway and methionine cycle. **B)** Western blot of A549, H2030, and H2009 NSCLC cell lines grown in High: 574.9 µM, Regular: 115.3 µM, and Low: 57.9 µM methionine RPMI media. Total H3 is used as a loading control. Protein ladder size is indicated on the left in kilo daltons (kDa). **C)** *CBS* mRNA expression in A549, H2030, and H2009 NSCLC cell lines in high, regular, and low methionine RPMI media, plotted as mean +/- SEM, n=4 biological replicates, ** indicates p<0.0028 by one-way ANOVA with Holm-Šídák’s multiple comparisons test. **D)** Western blot of A549 with and without LKB1 re-expression grown in high, regular, or low methionine RPMI media. Both β-actin and total H3 are loading controls. **E)** *LKB1* and *MAT2A* mRNA expression in A549 cells with and without LKB1 re-expression grown in high, regular, and low methionine media, plotted as mean +/- SEM, n= 4 biological replicates, ** indicates p=0.0013, ***p<0.0002, ****p<0.0001 by two-way ANOVA. **F)** RNA sequencing heatmap data of A549 cells with and without LKB1 re-expression grown in high, regular, and low methionine RPMI media. Genes involved in cell lineage, inflammation, and amino acid regulation were selected and grouped. **G)** Gene Set Enrichment Analysis of pathways involved in inflammation, cell cycle, and amino acid regulation using the RNA sequencing data of A549 cells with and without LKB1 re-expression in regular versus low methionine RPMI media. * indicates p<0.05 using the FDR q-value.

We next used RNA-sequencing to understand the global transcriptional consequences of differing methionine levels in both LKB1-mutant and LKB1-rescue A549 cells. In all levels of methionine, LKB1-rescue cells had down-regulation of bronchiolar cell markers, including *MUC5AC*, *SPEDF*, and *SOX2*, and an increase in transcripts that are associated with epithelial-mesenchymal transition, including *VIM*, *CXCR4* and *PDGFR* (**Figure 3F**). For inflammatory genes, the LKB1-mut cells had strikingly higher levels of cytokines that attract neutrophils, including *CXCL1*, *CXCL2* and *CXCL3*. Contrary to some other findings, it also appeared that LKB1- mut cells had higher levels of IFN-responsive genes, including *IFITM1* and *CD274* (encoding PD-L1), which could be explained by a lower level of the IFN inhibitor *MYC*. Together, these data fit our findings in mice that demonstrate that LKB1 loss leads to increased neutrophil infiltration and a subsequent transition to a more bronchiolar or squamous cell fate. We next queried genes in the methionine cycle (**Figure 3A**), and observed an intriguing pattern. Genes including the methionine transporters *SLC3A2* and *SLC7A5*, and the enzyme *MAT2A* were specifically up-regulated in the LKB1-mutant cells under low methionine conditions. Given the importance of MAT2A for SAM production, we confirmed the levels of this transcript by RT-qPCR (**Figure 3E**). Other genes regulated in this fashion included the genes involved in serine biogenesis, *PSAT1* and *PHGDH*. Finally, one of the most highly differentially expressed genes between the LKB1-mutant and LKB1-rescue cells, independent of methionine concentrations, was NNMT. NNMT serves as a ‘methyl sink’ and could further explain the observation that LKB1-mutant lung cancers have lower SAM levels [15, 24]. We next performed Gene Set Enrichment Analysis (GSEA [25]), and again found that pathways involved in amino acid deficiency and cell starvation were strongly included by low methionine conditions specifically in LKB1-mutant cells, but not in LKB1- rescue cells. LKB1-mutant cells also had stronger induction of ER-driven apoptosis, unfolded protein and protein translation responses, and larger increases in inflammatory responses in low methionine conditions relative to LKB1-rescue cells. For proliferation and PRC2 gene sets, both LKB1-mutant and -rescue cells had a depletion of cell cycle pathways in low methionine vs regular and high methionine, though the changes were most robust in the LKB1-deficient when comparing low to high methionine. Genes bound and repressed by the PRC2 member EED were increased in low methionine only in the LKB1-defienct cells, further supporting the loss of H3K27me3 in these cells (**Supp.** Figure 3B**+C**).

Finally, we wanted to further examine the relationship between LKB1 and CBS. Therefore, we developed additional transduced A549 cell lines with either CBS knock-down (shRNA) or CBS overexpression in both the control (LKB1-mutant) and rescued (LKB1-WT) cells. We found that knock-down of CBS could be achieved in both cell lines, and did not change LKB1 protein or RNA levels in the LKB1-rescue cells (**Figure 4A+B, Supp.** Figure 4A). When LKB1 was rescued, endogenous CBS levels were increased at both the mRNA and protein levels, and H3K27me3 were also increased, which could indicate that LKB1-rescue increased SAM pool levels (**Figure 4C+D, Supp.** Figure 4B). Strikingly, when CBS was expressed off a lentivirus in the LKB1-mutant cells, the levels of mRNA were increased but the protein levels were variable and relatively low. In contrast, in the LKB1-rescue cells, there was a clear and dramatic stabilization of both the mRNA and protein of the exogenous CBS, leading to an average of a 12-fold increase in protein expression. Together, these data suggest that LKB1 plays roles in regulating both CBS mRNA and protein levels, and that this regulation is unlikely to be through microRNAs which target the 3’UTR given that the exogenous construct only contained the open-reading frame.

**Figure 4:**
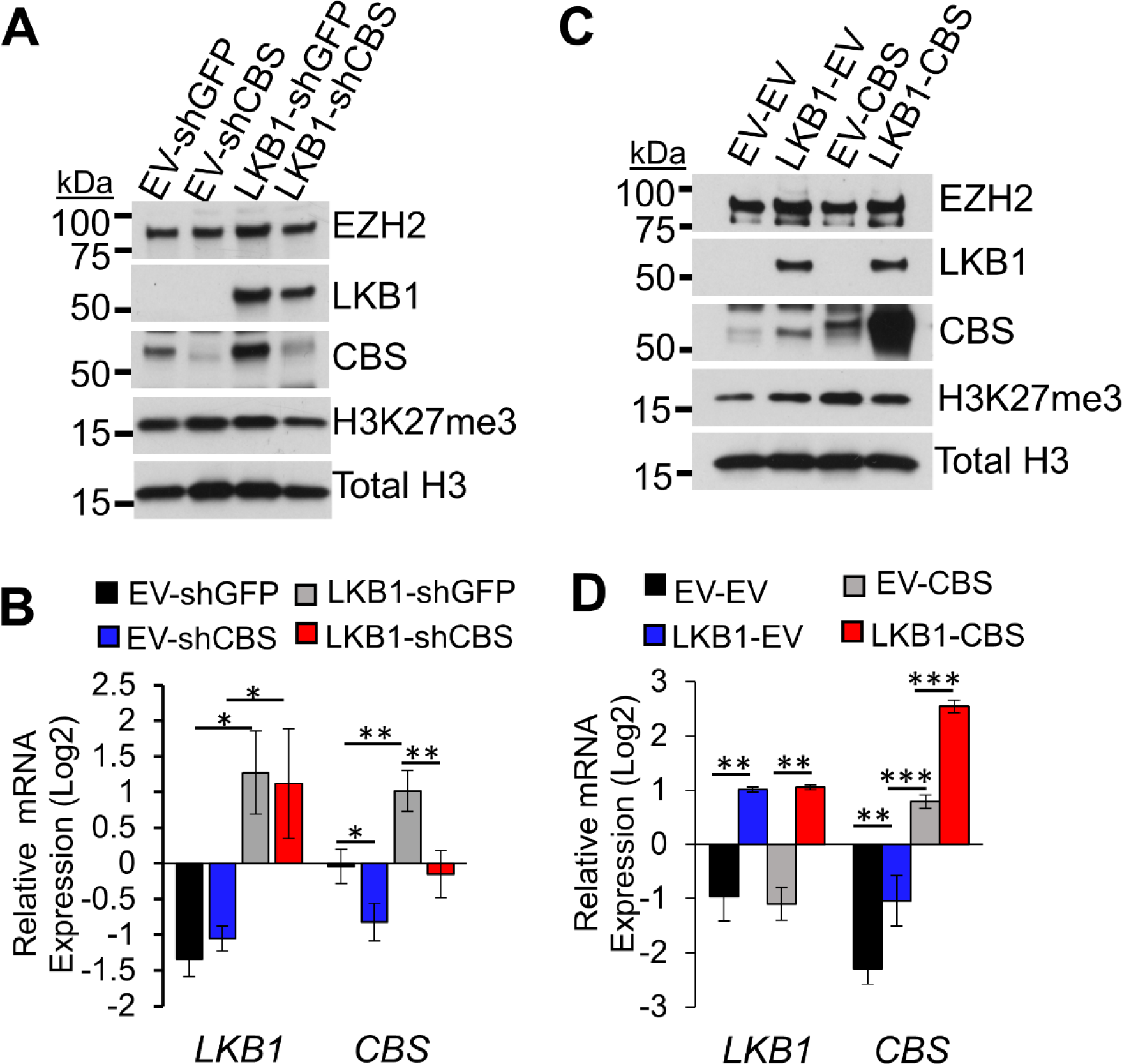
LKB1 Stabilizes CBS at mRNA and Protein Levels **A)** Western blot of A549 with and without LKB1 re-expression and with or without *CBS* shRNA knockdown. H3 is a loading control. Protein ladder markers are indicated on the left in kilodaltons (kDa). **B)** *LKB1* and *CBS* mRNA expression in A549 cells with and without LKB1 re-expression and with or without *CBS* shRNA knockdown, plotted as mean +/- SEM, n=8 biological replicates for *CBS* and n=4 biological replicates for *LKB1*, * indicates p<0.0298, **p<0.0032 by two-way ANOVA with Holm-Šídák’s multiple comparisons test. **C)** Western blot A549 with and without LKB1 re-expression and with or without CBS overexpression vector. H3 is a loading control. Protein ladder markers are indicated on the left in kilodaltons (kDa). **D)** *LKB1* and *CBS* mRNA expression in A549 cells with and without LKB1 re-expression and with or without CBS overexpression vector, plotted as mean +/- SEM, n=6 biological replicates for *CBS* and n=3 biological replicates for *LKB1*, ** indicates p<0.0059, ***p=0.0007 by two-way ANOVA with Holm-Šídák’s multiple comparisons test.

### LKB1 status and methionine concentration determines cellular responses to chemotherapy

Platinum-based chemotherapies remain one of the most common treatment strategies for advanced stage NSCLC [26]. To understand how LKB1 and CBS influence chemo responses, we pre-treated parental A549, H2030, and H2009 cells with high, regular, and low methionine media concentrations, and tested the responses to the common chemotherapy carboplatin. We found that carboplatin IC_50_ values were significantly lower in the LKB1-mutant A549 and H2030 cells grown in low methionine relative to regular methionine (**Figure 5A**). High methionine provided a strong carboplatin protective effect in both cell lines. In contrast, the LKB1-proficient H2009 cells had the same carboplatin IC_50_ values in both regular and low methionine. To examine these differences in a better controlled system, we again turned to the A549 LKB1-rescue cells. Comparing these lines, we observed that LKB1-rescue cells had a higher carboplatin IC_50_ value in regular methionine, and that this increased IC_50_ could be rescued by growth in low methionine media. Like the parental line, both cell lines had a strong decrease in carboplatin sensitivity (i.e. higher IC_50_) when grown in high methionine, and the lowest carboplatin IC_50_ values were observed in LKB1-mutant cells grown in low methionine, suggesting that carboplatin can be most effective in conjunction with MR when LKB1 is deficient in the tumor cells.

**Figure 5:**
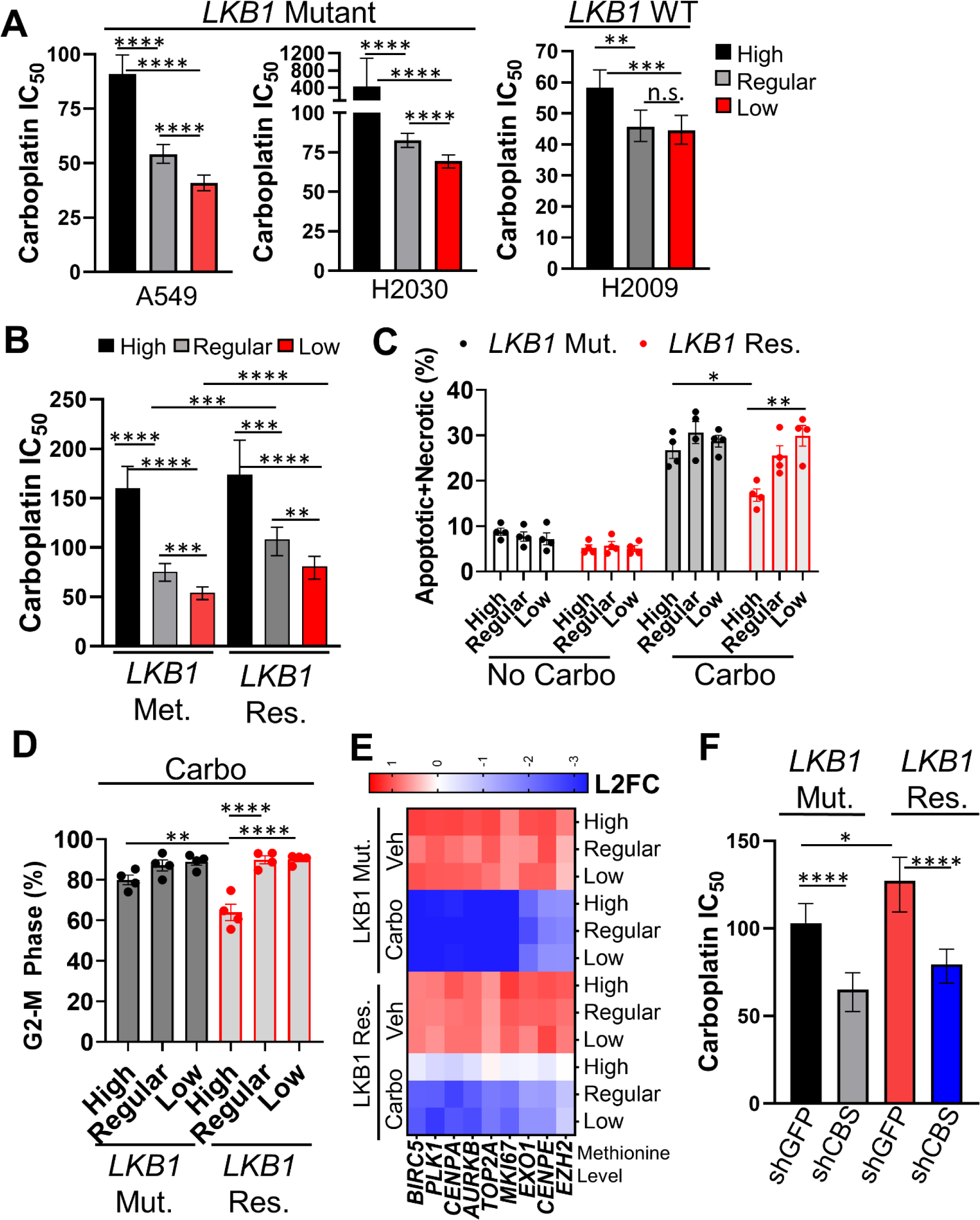
LKB1 Protects Cells From Carboplatin in a Methionine-Dependent Manner **A)** Carboplatin IC_50_ values in A549 (*LKB1* Mutant), H2030 (*LKB1* Mutant), and H2009 (*LKB1* WT) cell lines were obtained from non-linear regressions of carboplatin dose responses in high, regular, and low methionine RPMI media. Plotted as mean IC_50_ +/- the 95% confidence intervals, n=4 biological replicates for H2030 and H2009, n=5 biological replicates for A549. ** indicates p=0.0011, ***p=0.0002, ****p<0.0001 and n.s.=not significant by comparing IC_50_ values between the groups with sum of squares F test. **B)** Carboplatin IC_50_ values in A549 cells with and without LKB1 re-expression were obtained from non-linear regressions of carboplatin dose responses in high, regular, and low methionine RPMI media. Plotted as mean IC_50_ +/- the 95% confidence intervals, n=4 biological replicates. ** indicates p=0.0031, ***p<0.0004, and ****p<0.0001 by comparing IC_50_ values between the groups with sum of squares F test. **C)** Annexin-V apoptosis flow cytometry on A549 cells with and without LKB1 re-expression grown in high, regular, and low methionine media with and without carboplatin treatment at 60 µM, showing a combined percentage of apoptotic and necrotic cell population, plotted as mean +/- SEM, n=4 biological replicates. * indicates p=0.0242 and **p=0.0022 by one-way ANOVA with Holm-Šídák’s multiple comparisons test. **D)** 7AAD cell cycle flow cytometry on A549 cells with and without LKB1 re-expression grown in high, regular, and low methionine media with 60 µM carboplatin treatment, showing only changes to G2-M phase, plotted as mean +/- SEM, n=4 biological replicates, ** indicates p=0.0031, ****p<0.0001 by one-way ANOVA with Holm-Šídák’s multiple comparisons test. **E)** RNA sequencing heatmap data of A549 cells with and without LKB1 re-expression grown in high, regular, and low methionine RPMI media with and treated with 60 µM carboplatin. **F)** Carboplatin IC_50_ values in A549 cells with and without LKB1 re-expression and with and without CBS knockdown via shRNA were obtained from non-linear regressions of carboplatin dose responses. Plotted as mean IC_50_ +/- the 95% confidence intervals, n=4 biological replicates. * indicates p=0.015 and ****p<0.0001 by comparing IC_50_ values between the groups with sum of squares F test.

We next chose a carboplatin dose in around the regular IC_50_ of 60µM to perform apoptosis, cell cycle and RNA-seq analyses. Intriguingly, at this dose of carboplatin, the most dramatic differences were observed in the LKB1- rescue cells, which had less apoptosis (**Figure 5C**), less G2-M arrest (**Figure 5D, Supp.** Figure 5A), and a less dramatic decrease in cell cycle associated genes (**Figure 5E**). These results suggest that while high methionine is protective from carboplatin, LKB1-WT cells are the most able to exploit a high methionine environment to survive carboplatin treatment. Finally, to further investigate the effect of CBS expression on the response to carboplatin, we tested response of carboplatin in the CBS knock-down cells. Both the LKB1-mutant and LKB1- rescue cells CBS knockdown had significantly lower carboplatin IC_50_ values when compared to the shGFP control cells (**Figure 5F**). It has been shown that glutathione binds to platinum-based compounds, including carboplatin, and aids in transport of this xenobiotic from the cell. Given that CBS activity is upstream of glutathione production and is depleted by low methionine, these results suggest a mechanism through which CBS regulates carboplatin response.

### Methionine increases response to carboplatin chemotherapy *in vivo*

While the results in vitro suggested a role of MR in sensitization to carboplatin in LKB1-mutant cells, the true test of chemotherapy response is *in vivo*. To test response to carboplatin in a biologically relevant system, we induced tumors in the *Kras^mut^/*Lkb1^mut^ and separated cohorts into control chow and MR chow at 8 weeks post adeno-Cre. Mice were then further sub-divided to receive either carboplatin or sham once per week starting at 9 weeks post adeno-Cre, along with anti-nausea medications, for 3 cycles (**Figure 6A**). We analyzed the weights of these mice, and observed a steady decline in weight in all groups from week 9 to week 12, likely due to progression of disease and frequency of handling. Importantly, the MR chow did not lead to any additional weight loss when compared to control chow (**Supp.** Figure 6A). After the 3 cycles of chemotherapy, we inflated all five lobes of the lungs with formalin, sectioned, and analyzed the tumor characteristics (**Figure 6B**). For overall tumor burden, both carboplatin and MR were effective at reducing tumor burden, with the lowest tumor burdens in mice that received both treatments (**Figure 6A**). Furthermore, tumor number and the tumor burden of alveolar (A-ADC) tumors were statistically lower in carboplatin treated mice on MR than carboplatin treated mice on control chow (**Figure 6B+C**). Hyperplasia burden was also lowest in the MR/carbo group, suggesting that tumors of all stages are reduced by treatment. When comparing tumor burden of bronchiolar (B-ADC) the trends were similar, but due to variability among mice, the only statically significant difference was between control chow/sham and MR/carbo groups (**Supp.** Figure 6B). Taken together, these data indicate that MR can potentiate carboplatin response, even when started only a week prior to treatment, and that it is most effective at reducing alveolar adenocarcinoma tumors.

**Figure 6:**
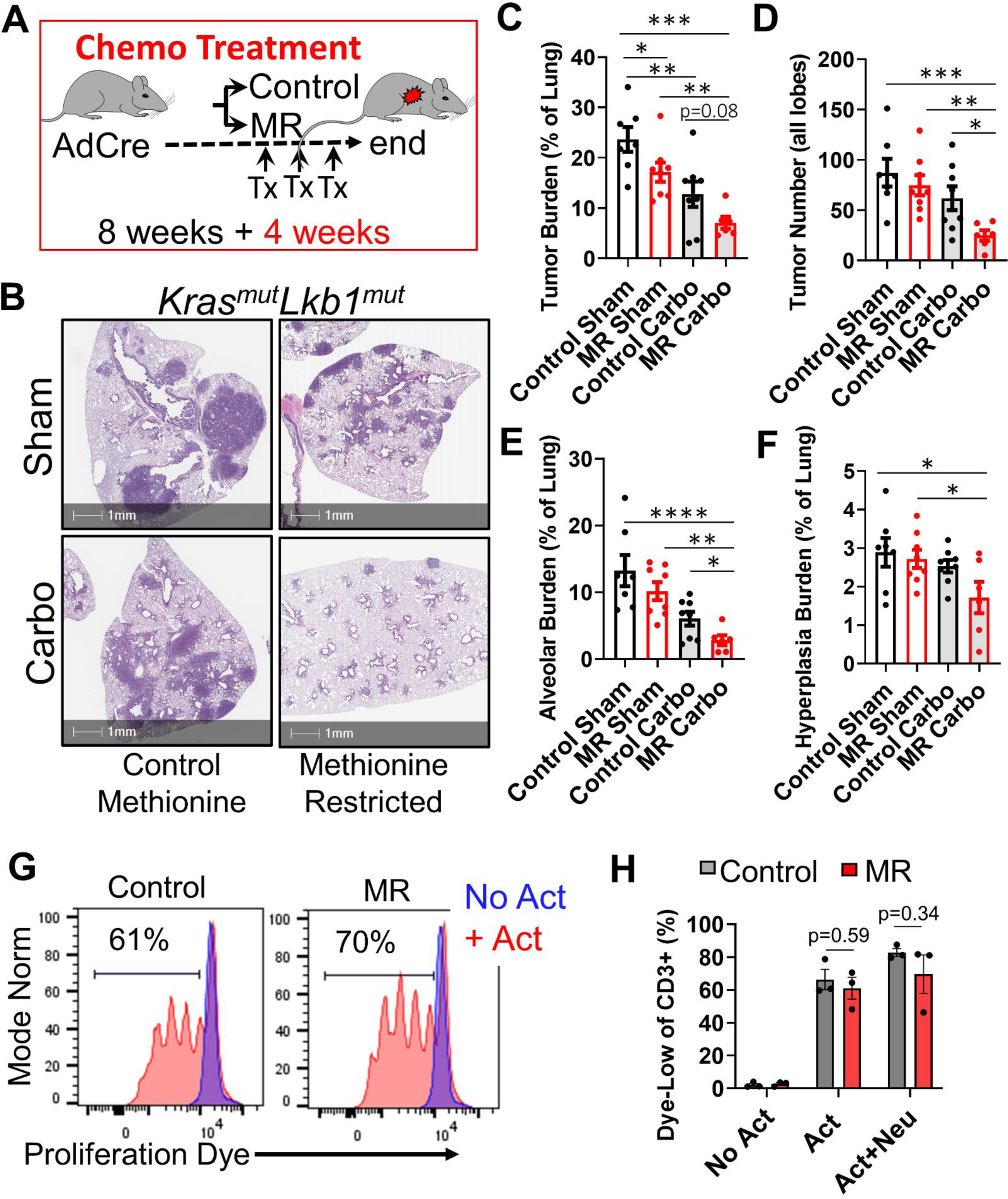
Carboplatin Response is Improved *in vivo* by Methionine Restriction **A)** Treatment schematic of methionine restriction and carboplatin chemotherapy. Mice were induced with Adeno-Cre virus via intranasal instillation. After 8 weeks, mice were separated into control and methionine restricted diet groups. One week post diet change, mice were given doses of anti-nausea (1µg/g maropitant and 0.8µg/g ondansetron) one hour prior to a dose of either sham treatment or carboplatin (60µg/g) via IP injections. Treatment was administered in this manner for 3 weeks (3 total treatments), and one week post the last treatment mice were sacrificed. **B)** Representative H+E staining images of lung lobes of mice on control and methionine restricted diets on both sham and carboplatin treatments. Scale bars=1mm. **C)** Tumor burden calculated as the percentage of the total lung area that was identified as tumor for mice on control and methionine restricted diets on sham and carboplatin treatments. Plotted as mean +/- SEM, n=7 for control sham, n=8 for control carbo, n=8 for MR sham, and n=6 for MR carbo. One mouse was removed from the MR carbo group after performing the ROUT outlier identification test, * indicates p=0.0414, **p<0.004, and ***p<0.0001 by one-way ANOVA, p value between control carbo and methionine restricted carbo is also indicated at p=0.08. **D)** Number of total tumors for mice on control and methionine restricted diets on sham and carboplatin treatments. Plotted as mean +/- SEM, n=7 for control sham, n=8 for control carbo, n=8 for MR sham, and n=6 for MR carbo, * indicates p=0.0301, **p=0.0048, and ***p=0.0009 by one-way ANOVA. **E)** Alveolar tumor burden calculated as the percentage of the total lung area that was identified as alveolar tumor for mice on control and methionine restricted diets on sham and carboplatin treatments. Plotted as mean +/- SEM, n=7 for control sham, n=8 for control carbo, n=8 for MR sham, and n=6 for MR carbo, * indicates p=0.0366 by two-tailed t-test, **p<0.0025, and ***p<0.0001 by one-way ANOVA. **F)** Hyperplasia burden calculated as the percentage of the total lung area that was identified as hyperplasia for mice on control and methionine restricted diets on sham and carboplatin treatments. Plotted as mean +/- SEM, * indicates p<0.0265 by one-way ANOVA. **G)** Representative cell proliferation dye flow cytometry of T-cells from both control and methionine restricted mice with and without T-cell activation by αCD3/αCD28. **H)** Proliferation dye quantified for three mice per group, mean +/- SEM is graphed, n=3 for all groups, p=0.34 and p=0.59 by one-way ANOVA with Holm-Šídák’s multiple comparisons test.

Due to reports that T cells require methionine to proliferate and that gut microbiome is influenced by MR, there is much debate as to whether MR can be successfully combined with immunotherapy [3, 27, 28]. For advanced lung cancer, carboplatin-based chemotherapy will be most often prescribed with anti-PD1/PD-L1 immunotherapy and therefore it is important to understand if MR can be combined with immunotherapy in this disease [26]. To understand if MR will impair T cells, we isolated splenocytes from tumor-bearing mice that had been on MR for 4 weeks. Due to their role as the effector cells for anti-PD/PD-L1, we used a kit to enrich for CD8 T cells as verified enrichment by flow cytometry (**Supp.** Figure 6C). We then stained the T cells with proliferation dye and plated them in wells pre-coated with CD3 and CD28 antibodies or wells with no activation antibodies as controls. T cells plated without activation antibodies were unable to proliferate and were used as gating controls for the flow cytometry (**Supp.** Figure 6D, **Figure 6D**). In wells with activation antibodies, T cells from MR mice, grown in low methionine conditions, proliferated as well as those from control chow mice grown in regular methionine conditions (**Figure 6D+E**). Due to a known role of neutrophils in T cell suppression, we also isolated neutrophils from bone marrow of the same mice and co-cultured these cells with the T cells. Addition of neutrophils did not prevent T cell proliferation in either MR or normal methionine conditions (**Figure 6E**). Together, these data suggest that T cells can undergo clonal expansion in low methionine conditions, and therefore, it may be possible to safely combine MR with immunotherapy for therapeutic intervention.

### CBS expression is highest in adenocarcinomas from smokers and LKB1-WT tumors

Finally, we sought to understand CBS levels in distinct NSCLC tumor subtypes. We first queried *CBS* expression in the TCGA lung adenocarcinoma and lung squamous cell carcinoma datasets. We separated the samples into normal (tumor adjacent) and into groups based on smoking status. We observed that *CBS* transcript levels were low in ADC tumor-adjacent normal tissue, and highest in ADC tumors from patients that were current smokers (**Figure 7A**). *CBS* levels were lower in ADC tumors from patients that were reformed or never smokers, which suggest that high levels in smokers are due to smoking-induced oxidative stress. In SCC, *CBS* levels were also higher in tumors than in tumor-adjacent normal tissue, but levels were not different among groups with different smoking status. EZH2 levels followed similar trends, although as seen previously [8, 9], SCC have higher levels of EZH2 and again levels were not changed by smoking (**Supp.** Figure 7A). Finally, we examined levels of glutathione synthase (*GSS*) (**Figure 7B**). GSS levels were higher in SCC and its adjacent tissues than in ADC, independent of smoking status. Next, due to regulation of CBS at the protein level, we tested CBS protein expression in a tissue microarray that contained adenocarcinoma, squamous cell carcinoma and adenosquamous tumors. We further subdivided the adenocarcinomas into alveolar (A-ADC), bronchiolar (B- ADC) and solid type, poorly differentiated tumors. Using HALO, we measured CBS staining in both the tumor (epithelial cell) and stromal (mesenchymal and immune cell) compartments (**Figure 7C**). We observed that A-ADC/B-ADC and adenosquamous type tumors had high levels of CBS, while solid ADC and squamous cell carcinomas had lower CBS levels, and in all tumor types CBS was higher in the malignant epithelium compared to the cells in the stromal compartment (**Figure 7D, Supp.** Figure 7B). It has been postulated that oxidative stress helps drive transition of adenocarcinomas into adenosquamous and squamous phenotypes, and these data support the idea that CBS is highest in tumors with high oxidative stress levels, which are then relieved by full transition into the squamous state. Together these data suggest relationships between CBS levels and mutational status and lineage of lung cancers.

**Figure 7:**
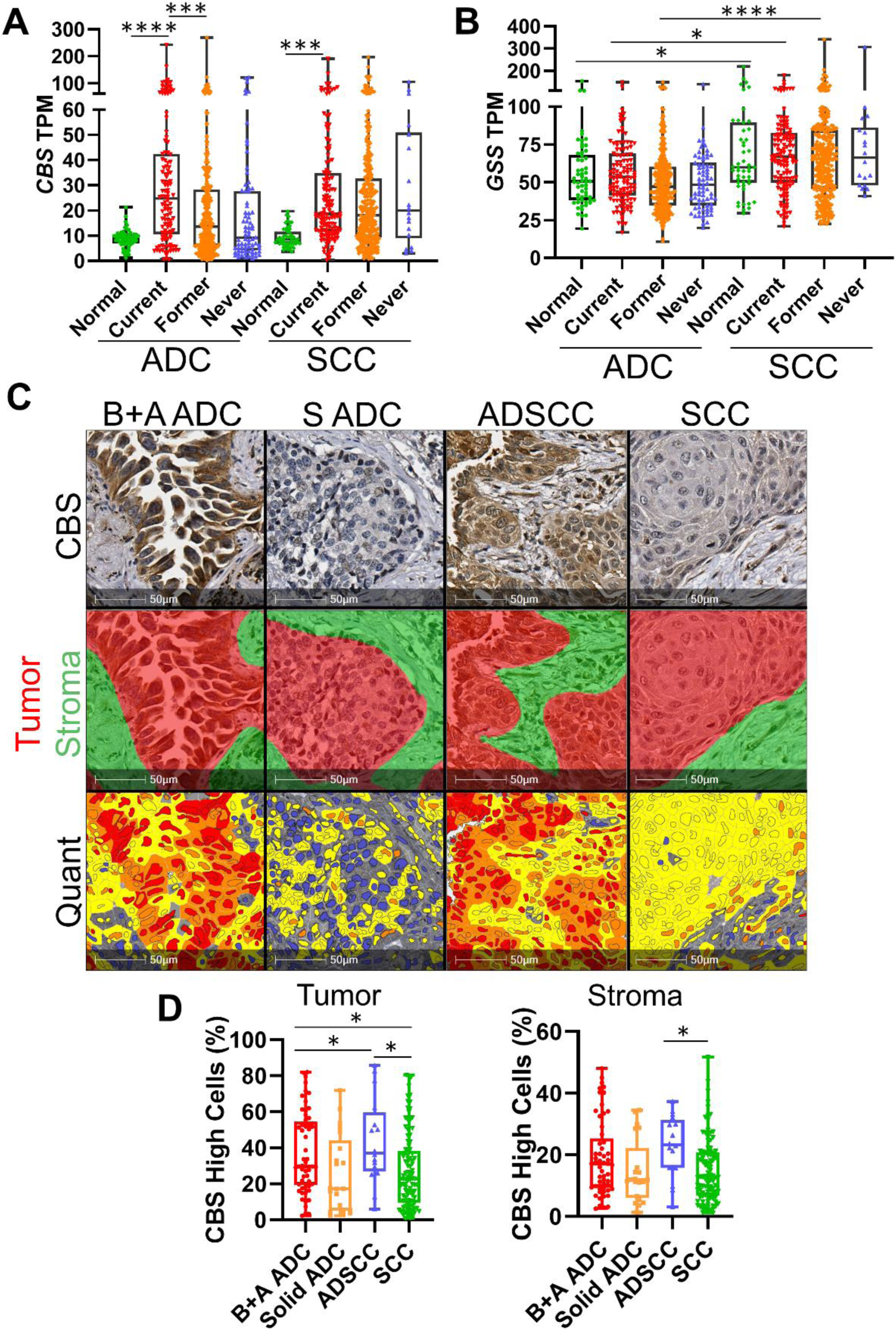
CBS is Highest in Smokers, Adenocarcinomas and Adedosquamous Lung Tumors **A)** Human TCGA data from patient samples grouped by tumor subtype and smoking status for *CBS* expression (left panel) and *GSS* expression (right panel) in transcripts per million (TPM). Groups are Normal A (Normal adjacent Adenocarcinoma, n=58), Current A (Current smoker Adenocarcinoma, n=120), Former A (Former smoker Adenocarcinoma, n=307), Never A (Never smoker Adenocarcinoma, n=75), Normal S (Normal adjacent Squamous Cell Carcinoma, n=51), Current S (Current smoker Squamous Cell Carcinoma, n=133), Former S (Former smoker Squamous Cell Carcinoma, n=338), and Never S (Never smoker Squamous Cell Carcinoma, n=18), * indicates p<0.021, ***p=0.0001, and ****p<0.0001 by one-way ANOVA with Holm-Šídák’s multiple comparisons test. **C)** Representative immunohistochemistry for CBS with HALO tumor vs stroma and quantification algorithms, scale bar = 100µm. **D)** Quantification of indicated markers in diverse lung tumor types, plotted as box-and-whisker plots, error bars are min-max, box bounds are 25th and 75th percentiles and center line is median, *p<0.0495 by one-way ANOVA with Holm-Šídák’s multiple comparisons test.

## DISCUSSION

Here, we demonstrate that methionine restriction (MR) diet is an effective therapeutic strategy for KRAS/LKB1 mutant NSCLCs. Using MR in this model resulted in decreased tumor burden, tumor size, and hyperplasia burden. Furthermore, tumors in MR mice were less likely to progress into a squamous state, suggesting that MR prevented the progression of these tumors. Finally, we analyzed the TIMEs and found that neutrophils, a cell type implicated in the transition from adenocarcinoma to squamous cell carcinoma [19, 21, 29], were significantly less abundant in tumors from MR mice relative to those in control chow mice. Previous studies have shown that LKB1 loss leads to a rapid accumulation of neutrophils that accompany [8, 30], and perhaps drive [29], the adeno-squamous transdifferention in the *Kras^mut^/*Lkb1^mut^ mouse model. Intriguingly, we observed that LKB1 rescue in human lung cancer cells led to decreased expression of neutrophil attracting cytokines *CXCR1/2/3* and the squamous transcription factor *SOX2*, and increased global H3K27me3. These results mirror our previous findings that H3K27me3 is lost at *Sox2* and *Cxcl3/5/7* genomic loci during squamous transition, and that *Lkb1* deletion provides a strong induction of these cytokines and influx of tumor-associated neutrophils [8]. These data indicate that at least in the A549 cell line, observations from the mouse model hold in human cells, demonstrating that loss of PRC2 is a likely driver of adeno-squamous transition.

LKB1 is essential for regulation of energy consumption, and deficiencies in LKB1 have been shown to disrupt the metabolic states of cancer cells in many ways, including lowering levels of SAM pools *in vivo* [10, 15, 31]. In KRAS/LKB1 mutant cells, H3K27me3 was reduced by MR, which was in contrast to a rather specific reduction in H3K4me3 observed in MR colorectal tumors [32]. Our observation is supported by the fact that SAM levels can be significantly increased by an EZH2 inhibitor in KRAS/LKB1 mutant cells [33], suggesting that PRC2 is a major utilizer of SAM in this genotype of cells. It is important to note that this same paper showed that LKB1 rescue in H1944 and H2122 cell lines lead to *decreased* SAM levels, indicating that other genetic determinants may control this phenotype, or that total SAM levels in cells do not directly correlate to nuclear SAM availability [33]. We also found that cells deficient in LKB1 grown in low methionine media had increases in inflammatory response genes, and a dramatic increase in *MAT2A* expression. It is known that *MAT2A* transcript is degraded in response to high SAM levels [34, 35], further implying that SAM availability drops below a homeostatic threshold in LKB1-mut cells grown in low methionine.

Furthermore, LKB1 status influenced the response of cells to chemotherapy in a methionine dependent fashion. LKB1-WT cells were provided a protective effect against carboplatin treatment in all methionine concentrations relative to LKB1-mut cells. Intriguingly, LKB1-WT cells also had less G2M arrest and less apoptosis when grown in high methionine media, relative to responses of LKB1-mut cells. To validate these findings, we demonstrated that MR can be coupled with chemotherapy treatment to increase response *in vivo*. We also observed that LKB1-WT tumor cells had higher levels of the transsulfuration pathway enzyme CBS than those with mutant LKB1. Knock-down and overexpression experiments demonstrated that LKB1 and CBS expression are linked, and that CBS expression is partially dependent on LKB1 status. The link between LKB1 mutation and CBS expression has not been observed previously. However, SAM is known to activate and stabilize CBS, further suggesting that SAM is more abundant in LKB1-WT cells [36]. Furthermore, we found that knock-down of CBS could sensitize cells to carboplatin, both in LKB1-mut and LKB1-WT cells. The most likely explanation for this result is decreased glutathione levels. Glutathione has been shown to bind platinum-based compounds like carboplatin and aid in transport of these xenobiotics from the cell [37]. Furthermore, CBS was recently found to be a poor prognostic biomarker in squamous cell carcinomas, and this may be due to an enhanced chemotherapy resistance of these tumors [38].

Because immunotherapy is quickly becoming common treatment for nearly all NSCLC patients, understanding how MR influences T cells is an important question. Previous reports have been conflicting on the roles of methionine metabolism in T cells and immune responses. For example, a recent study found that SAM and its metabolite MTA lead to T cell exhaustion [39]. However, several studies have shown that methionine is required for proper T cell differentiation [27, 40], and that MR alters the gut microbiome to cause immunotherapy resistance [28]. Our data indicate that T cells from MR tumor-bearing mice can be expanded in low methionine media just as efficiently as those from control chow mice. These results suggest that MR can be combined with immunotherapy for lung cancer, a theory that our lab will test next. One potential challenge of implementing MR diets in the clinic is that the diet is essentially vegan [41], and patients may not want or be able to follow a diet that prevents them from enjoying certain foods. However, while our data indicates there are benefits to MR long-term, there are also short-term benefits. We were able to show that after only 4 weeks on a MR diet, our mice had reduced tumor burden compared to the control methionine mice, and that MR was able to cooperate with chemotherapy in a short time frame. This suggests that patients may only need to adhere to stricter diets prior to receiving, during, and shortly after treatments. Recent data also suggests that intermittent MR, decreasing methionine intake a few days a week, could impart significant benefits to immunotherapy-based treatment [42]. Knowing this, designing trials where patients switch their diet a few days a week may increase the likelihood that patients would adhere to these strict diets.

Overall, we have provided evidence that MR would benefit patients as a therapeutic intervention for aggressive KRAS-driven lung cancers. While more research is needed into how MR could impact the TIME, our initial results provide promising groundwork. In addition, we show evidence for the roles of MR, LKB1, and CBS expression in the lineage fate of NSCLCs. These data pave the way to formulate more research with MR with the end goal of improving lung cancer patient outcomes.

## EXPERIMENTAL PROCEDURES

### Mouse Cohorts

Animal work in this study was approved by the University of Kentucky following regulations outlined by the Institutional Animal Care and Use Committees (IACUCs). All animal work in this study was conducted using a genetically engineered mouse model (GEMM) of non-small cell lung cancer containing a single *KrasG12D* allele [43] with loxP sites flanking a STOP codon and both alleles of *Lkb1* flanked by loxP sites [44] (LSL: *KrasG12D*; *Lkb1* flox/flox) leading the conditional *Kras^mut^/*Lkb1^mut^ mice [18]. Mice were maintained on Teklad chow 2918 (methionine 0.4% w/w) until randomization to treatment arms. Adult mice were administered adeno-CMV-Cre virus at a final concentration of 3 x 10^7^ PFU (University of Iowa) per mouse via intranasal instillation. Once administered, Cre-recombinase removes the STOP cassette flanking the *Kras* gene, activating the *KrasG12D* mutation, and deletes a significant portion of the *Lkb1* gene, rendering it inactive. Genotyping primers are in **Supplementary Table 1**.

### Methionine Restriction for Tumor Prevention

For the tumor prevention study, one week post administration mice were split into two groups: control methionine chow at 0.86% w/w methionine and low methionine chow at 0.12% w/w methionine (LabDiets.Com A11051301B and A11051302B, respectively). A total of 10 males and 10 females were placed into each diet category. Mice were monitored twice a week for weight loss. 10 weeks post administration (9 weeks post diet change), mice were sacrificed. Blood was collected via cardiac punctures, and serum was isolated from blood via centrifugation at 14,000 rpm and 4°C for 15 minutes. Metabolites in the serum were stabilized with the addition of 4.5 µL of formic acid/mL of serum. Stabilized serum was then flash frozen in liquid nitrogen and stored in -80°C until ready to use. Lungs were inflated with 10% formalin and then collected for paraffin embedding and sectioning. Several sections were cut, including sections for Hematoxylin and Eosin (H&E) staining and immunohistochemistry (IHC). H&E stains were analyzed on the HALO imaging software to calculate tumor burden, number and size as well as examining immune cell infiltration with our published nuclear phenotyper algorithm [22].

### Methionine Restriction Coupled with Chemotherapy

For the chemotherapy treatment study, 8 weeks post administration of adeno-CMV-Cre, mice were separated into the same two dietary groups of control (0.86% w/w) and low (0.12% w/w) methionine chow. Mice were sensitized to the diet for 1 week. After 1 week, mice were first treated with a mixture of 1µg/g maropitant and 0.8µg/g ondansetron for anti-nausea via subcutaneous injections. One hour after anti-nausea injections, 60µg/g carboplatin (Millipore) in saline or placebo (saline only) was administered via IP injections. The following two days (24 and 48 hours post carboplatin), mice were given an additional dose of anti-nausea medication at the same concentrations. Anti-nausea and carboplatin doses were administered in this manner for 3 weeks. One week post the last dose of carboplatin, the mice were sacrificed. Lungs and serum were collected and calculations for tumor and hyperplasia burden were obtained in a similar manner to the prevention study. In one cohort several mice had no visible tumors, likely due to issues with intranasal adenovirus administration, and those mice were not included in the final analysis.

### Human Lung Cancer Cells

All cell lines were maintained following University of Kentucky biosafety guidelines. Human non-small cell cancer A549, H2009 and H2030 cells were obtained from ATCC. Cells were cultured in RPMI 1640 media (Gibco, CAT#: 11875-093) supplemented with 9% FBS (VWR, CAT#: 97068-0085), 0.9x Penicillin-Streptomycin (Gibco, CAT#: 15140-122) and 1x GlutaMAX^TM^ (Gibco, CAT#35050-061) (unless indicated otherwise) in 37°C, 5% CO_2_ and 21% O_2_. Cells were treated with Plasmocin (InvivoGen, CAT#: ant-mpt) after thaw for about 1 week and tested for mycoplasma regularly using the MycoAlert^TM^ PLUS Mycoplasma Detection Kit (Lonza, CAT#: LT07-318). Cells were always used within 10 passages of authentication via STR analysis with CellCheck9 by IDEXX laboratories.

### Cloning of Viral Vectors and Transduction of Cells

A pBABE-puro vector (a gift from Hartmut Land & Jay Morgenstern & Bob Weinberg, plasmid #1764 on AddGene) [45] and pBABE-FLAG-LKB1 vector (a gift from Lewis Cantly, plasmid #8592 on AddGene) [14] in bacteria were ordered from AddGene as bacterial stab stocks. A single bacteria colony was selected from bacteria grown on agar plates (1% Tryptone, 0.5% Yeast Extract, 1% NaCl, 1.5% agar) after expansion in Luria Broth (1% Tryptone, 0.5% Yeast Extract, 1% NaCl) both supplemented with 0.1 mg/mL ampicillin. This single colony was expanded and plasmid was isolated utilizing the Qiagen QIAprep Spin Miniprep Kit (CAT#: 27106). CBS overexpression and control vectors were made previously [46]. shCBS knockdown vectors were prepared using CBS primers obtained from Integrated DNA technologies (Forward: 5’ – CCG GGC GGA ACT ACA TGA CCA AGT TCT CGA GAA CTT GGT CAT GTA GTT CCG CTT TTT G – 3’; Reverse: 5’ – AAT TCA AAA AGC GGA ACT ACA TGA CCA AGT TCT CGA GAA CTT GGT CAT GTA GTT CCG – 3’) and annealed and ligated in the pLKO.1 Hygro linearized backbone (a gift from Bob Weinberg, plasmid #24150 on Addgene). Stbl3 bacteria were heat-shocked to uptake ligated plasmid, expanded in Luria broth, and single colonies on an agar plate (both broth and agar supplemented with 0.1 mg/mL ampicillin) were selected and sequenced for inserts. Control shGFP in the pLKO.1-Hygro vector were made previously [46]. The KRAS, LKB1-rescue, shCBS, and CBS OE vectors and each corresponding control lenti- or retro-viral vectors were transfected into HEK293T cells (lenti) or PlateE cells (retro) using VSVG, vprΔ8.2 for the cells to produce live virus for infection. Virus-containing media was collected from packaging cells and stored at -80°C for future use. For transduction, A549 or BEAS2B cells were plated at low confluency in 6-well plates for infection. Cells were allowed to adhere to the plate before continuing (usually overnight). Once adhered, 1.5 mL of fresh media was added to each well and 500 µL of viral vector was added to each well. Cells were placed back in the incubator overnight. Virus was removed the next morning and replaced with fresh media. Cells were allowed to grow for the several days, then split to 10 cm dishes for selection with 1µg/mL puromycin (pBABE/LKB1 vectors), 10 µg/mL Blasticidin (EV/CBS overexpression vectors), and 0.2 mg/mL Hygromycin (shGFP/shCBS vectors), or sorted for GFP expression. Selected cells were used for further experiments described below.

### Growth of Human Cell Lines in Methionine Modulated Media

A549, H2009, H2030, BEAS2B, and A549 cells with different transduced with lentivirus (pBABE/LKB1, EV/CBS, shGFP/shCBS) were grown and maintained in regular RPMI media (see Human Lung Cancer Cells Method). To run experiments with different concentrations of methionine, cells were plated in 10 cm dishes at 200k cells/plate and sensitized to high methionine, regular methionine and low methionine media RPMI media. Methionine media was prepared using either methionine-free RPMI media (Invitrogen) or DME-F12 SILAC media (AthenaES, CAT#: 0432). All media was supplemented with FBS, Penicillin/Steptomycin, and Glutamax in the same concentrations in the regular RPMI media. DME-F12 was also supplemented with leucine, lysine and arginine to base media level concentrations and insulin/transferrin/selenium supplement was added. Media was then separated into 3 equal aliquots and supplemented with different concentrations of methionine (High: 574.9 µM, Regular: 115.3 µM, Low: 57.9 µM). Cells were sensitized to these concentrations of methionine for 5 days and fed with fresh media on day 3. On day 5, cells were collected and/or counted for various experiments.

### Carboplatin Dose Response Assays

For the dose response assay, cells were plated into black opaque 96-well plates at 2,500 cells/well in the same concentration of methionine media they were sensitized to and allowed to adhere to the plate for 1 day. The next day, cells were fed 50 µL of the same methionine media with 10 concentrations of carboplatin between 0-125 µM (SIGMA). On day 4, 50 µL of Cell Titer Glo (Promega) was added to each well in the 96 well plate. Plates were mixed briefly on a shaker and then luminescence was measured on a Cytation5 machine. A non-linear regression of the data was created and used to analyze the IC_50_ for each cell line for carboplatin. For the parental A549, H2009, and H2030 cells, this experiment was completed in both RPMI and DME-F12 SILAC medias with the same results, and results from RPMI are shown.

### AV Apoptosis Flow Cytometry

For the apoptosis assay, A549 cells transfected with either pBABE-puro or pBABE-LKB1-FLAG vectors were plated into 10 cm dishes at 200k cells/plate in the same concentration of methionine media they were sensitized and allowed to adhere for 1 day. The next day, cells were treated with 60 µM carboplatin, which was determined based on the IC_50_ values for the cells on different concentrations of methionine. Media was not changed throughout the 4-day experiment. On day 4, media from the plates was collected and cells were collected with trypsin. Cells were counted and 200k cells were used for apoptosis. Cells were resuspended in 100 µL of Annexin binding buffer (BioLegend, CAT#: 79998). Cells for controls (from all samples) were mixed and resuspended in 300 µL of binding buffer and divided into tubes for double positive stain, 7AAD single stain, and unstained sample. 5 µL of FITC antibody (BioLegend, CAT#: 640906) and 5 µL of 7AAD (Invitrogen, CAT#: A1310) were added to each sample and to the double positive tube. Only 7AAD was added to the 7AAD single stain tube, nothing was added to the unstained tube. Cells were resuspended in 400 µL of Annexin binding buffer and then filtered through a FACS tube in a 35µm filter. Samples were run through a BD Symphony flow cytometer. Data were analyzed using FloJo and precent apoptotic, necrotic and live cells were compared for all samples.

### Cell Cycle Flow Cytometry

The same cells collected for the apoptosis assay were used for the cell cycle assay. 200k cells were placed in a 1.5 mL tube and spun down. Single stains were prepared in similar fashion as above in *Apoptosis* for an unstained and a 7AAD tube. Media was aspirated and cells were resuspended in 300 µL of PF10 (10% FBS in PBS). 700 µL of 70% ethanol was added slowly to each tube and vortexed to mix. Tubes were left to fix at 4°C at least overnight. After fixing, cells were resuspended in 100 µL of 1 mg/mL RNaseA (Thermo Scientific, CAT#: EN0531) in PBS and allowed to sit for 30 minutes at room temperature. Cells were then washed with PBS. A solution of 12.5µg/mL 7AAD (Invitrogen, CAT#: A1310) antibody to PBS was made. Cells were resuspended in 250 µL of this solution. The unstained tube was resuspended in 250 µL of just PBS. Samples were filtered through a FACS tube in a 35µm filter. Samples were run through a BD Symphony flow cytometer. Data were analyzed using ModFit LT to determine the percentage of cells in the G1, S and G2 phases of the cell cycle.

### RNA Isolation, RT-qPCR and RNA-sequencing

Cells were collected via trypsinizing and were lysed with RNA lysis buffer, made by mixing 100 µL of lysis buffer from the Agilent RNA isolation kit (CAT#: 400805) with 0.7 µL of β-mercaptoethanol (BME) per sample of cells. Cells were resuspended in this buffer. At this point samples were either further isolated or placed in -80°C until further use, extra RNA after making cDNA is also stored in the -80°C. RNA was isolated by following the microprep RNA isolation protocol and kit by Agilent (CAT#: 400805). RNA concentration after elution was calculated using a nanodrop in ng/µL of RNA. 1000 ng of RNA was used to make cDNA and mixed with 2 µL of a 1:1 mixture of random hexamers (50 ng/µL) and dNTPs (10 mM) and nuclease free water was added up to a total volume of 10 µL. PCR tubes were placed in the thermocycler and heated to 65°C for 5 minutes. After 5 minutes, the thermocycler is programed to go to 4°C, samples were allowed to sit at 4°C for 2 minutes before proceeding. Then, 10 µL of a mixture of 2x RT buffer, 10 mM MgCl2, 20 mM DTT, 2 U/µL RNase Out (Invitrogen) and 10 U/µL SuperScript III (Invitrogen) was added to each sample. Samples were placed back in the thermocycler and allowed to finish the cDNA protocol. After cDNA synthesis, 1 µL of RNaseH was added and incubated at 37°C for 20 minutes. Final cDNA was diluted 1:5 with nuclease free water. Sample reactions multiplexing gene of interest (FAM) with housekeeping probe (VIC) using Taqman reagents were assembled in 96 well plates and amplified in the StepOne Plus machine using the standard fast Taqman protocol. For RNA sequencing 1000ng of RNA was diluted in nuclease free water and sent to Innomics for sequencing using DNBseq at 150bp PE reads and 20 million reads per sample. To remove adapters and low-quality bases, reads were trimmed and filtered using Trimmomatic (V0.39) [47], and subsequently, reads were mapped to Ensembl GRCh37 transcript annotation (release 75) using RSEM [48]. Gene expression data normalization and differential expression analysis were performed using the R package edgeR [49].

### Gene Set Enrichment Analysis

GSEA was performed with GSEA version 4.3.2 (Broad Institute) with rank-ordered gene lists generated using all log-fold change values. Databases queried included Hallmarks (h.all.v2023.2), Reactome (c2.cp.reactome.2023.2), GO BP (c5.go.bp.v2023.2), and Oncogenic Signatures (c6.all.v2023.2). Gene signatures that converged on common themes were selected for further analysis and graphing of enrichment scores and FDRs for specific contrasts.

### Soft Agar Colony Formation Assay

BEAS-2B normal human lung cells were grown on plates coated with 0.1% gelatin (all BEAS-2B cells cultured in 2D format will be seeded in plates coated with gelatin unless otherwise stated) in normal RPMI media in 10 cm plates. Once confluent, 200k cells were seeded in 10 cm plates in High, Regular, and Low methionine RPMI media. Cells grew for 1 week sensitizing to the media. After 1 week, 20k cells were seeded in High, Regular, and Low methionine RPMI media and infected with KRAS-G12V virus [17] the following day. One week later, cells were sorted for GFP expression by FACS and 200k were seeded in High, Regular, and Low methionine media. After 5 days, cells were seeded in soft agar containing High, Regular, and Low methionine RPMI media. A 1.2% agarose solution and a 0.8% low melting agarose solution were prepared in PBS and autoclaved. Low melting agarose is kept in 37°C water bath to prevent solidifying. A mixture of 1:1 warmed media to 1.2% agarose solution was made and mixed thoroughly. 4 mL of this solution was added to each well in a 6-well plate and allowed to solidify at 4°C for about 2 hours. Volume of cells needed were mixed with 2 mL of 0.8% low melting agarose solution and 2 mL of warmed media and added on top of the already solidified agarose in each well and allowed to solidify t 4°C for 30 minutes. Before placing in tissue culture incubator, 3 mL of media was added to the top of each well, and 0.5 mL of media was added twice a week. Cells originally in low methionine media were seeded in soft agar containing all 3 methionine concentrations, taking total number of cell lines from 3 to 9 once seeded in the soft agar. Cells were allowed to grow in the soft agar for 1 month, they were then fixed with 100% methanol for 15 minutes. Colonies were then imaged and counted.

### Methionine and SAM Concentration from Murine Serum

Flash frozen serum stabilized with formic acid was thawed on ice. Bridge-IT SAM and Bridge-IT methionine kits from Mediomics were used for determining methionine and SAM concentrations in serum. Serum was pulse spun in a centrifuge (pulse button held until centrifuge reached 14,000 rpm) to collect any particulates to the bottom of the tube. Samples of serum were then used to run the assays as described in the assay protocols provided by the manufacturer. Standard curves for both methionine and SAM were created using the kit and manufacturer-provided protocols and then used to determine the concentration of methionine and SAM in the serum samples. After using the standard curve to calculate concentrations of each metabolite, dilution factors were used to determine actual concentration of the pure serum sample based on the dilutions used.

### Immunohistochemistry

Unstained paraffin wax slides were used to perform IHC staining for CBS. Slides were de-waxed and citrate buffer antigen retrieval was performed with citrate buffer. Tissues were circled with wax pen, and incubated with 10% goat serum in PBST for 45 minutes at room temperature in a humid chamber box. The goat serum was removed from the slides by gently tapping them on paper towels. Primary antibodies were diluted in 10% goat serum in PBST (CBS 1:200 Novus) and placed on the slides. Slides were incubated in a humid chamber at 4°C overnight. Primary antibody was removed from the slides by placing them in PBS in a slide holder and washing 3x for 5 minutes on a slide shaker. Slides were then kept in PBS prior to incubating with secondary antibody. Secondary antibody was prepared using Vector Labs ABC kit and the manufacturers protocol. DAB chromogen (Vector Labs) was used to visualize signal and Harris’s hematoxylin was used as counter-stain.

### Western Blotting

2D human non-small cell lung cancer cells were plated at 200k cells/10 cm dish and grown for 5 days until close to 100% confluency. Cells were washed with ice cold PBS (PBS chilled to 4°C) and collected by scraping in ice cold PBS. Supernatant was removed and cells were then flash frozen in liquid nitrogen and stored in -80°C until further use. Cell pellets were thawed on ice when ready to isolate protein. Protease and phosphatase inhibitor (1x Halt Buffer, Invitrogen) was added to RIPA buffer (50mM Tris, 150mM NaCl, 0.5% Na-Deoxycholate, 1% IGEPAL-CA630, 0.1% SDS) and mixed. About 50 µL of this lysis buffer (or more if a larger pellet) was added to the cell pellets and lysed via aggressive pipetting for 2 minutes. Cell lysate was then flash froze in liquid nitrogen and then placed back on ice. This was repeated several times over the course of 1 hour and 30 minutes. Cell lysates were always kept on ice and stored at -80°C. Lysate concentrations were calculated through a BCA assay and equal amounts of protein (50µg-1µg) were added to Laemmli loading dye, boiled for 5 minutes and placed on ice to cool. 4-12% acrylamide gels (BioRad) were loaded with protein samples. Gels were run at 100V for about 1.5 hours at RT. Protein was then transferred to nitrocellulose membranes using glycine transfer buffer on ice for 1 hour at 100V. Protein ladders on membranes were marked with a pen. Membranes were then blocked with 5% BSA in TBST for 1 hour at RT. Once blocked, membranes were cut for different antibodies depending on protein size and placed into the appropriate antibody solution made in TBST (CBS 1:1000 Cell Signaling Technologies CAT#: 14782S, EZH2 1:250-1:500 Cell Signaling Technologies CAT#: 5246S, LKB1 1:1000 Cell Signaling Technologies CAT#: 3047S, H3 1:5000 Abcam CAT#: ab1791, H3K27me3 1:500-1:1000 Cell Signaling Technologies CAT#: 9733S, ß-actin Abcam CAT#: ab8226). Membranes were stored at 4°C overnight while shaking on primary antibody. The next day, membranes were washed 3x with TBST for 5 minutes to remove primary antibody and then incubated at RT for 1 hour with secondary antibody (mouse Novus CAT#: NB7539, and rabbit Novus CAT#: NB7160) diluted 1:10,000 in TBST. They were then washed for 20 minutes 3x in TBST at RT. H3 strips were subject to additional washing if signal was too dark upon first exposure. Signal was activated using the SuperSignal West Pico PLUS Chemiluminescent Substrate (Thermo Scientific, CAT#: 34580). Excess luminescence solution was removed from membranes, and they were placed into an exposure chamber and exposed onto X-ray film (Amersham) in a dark room for several different exposure times (1 sec -2 minutes depending on signal strength of different antibodies). We quantified western blots using the ImageJ Gel Analyzer tool on 16-bit, inverted grayscale images.

### T-Cell Isolation/ Proliferation Assay

T-cells were isolated from the spleens of the mice by crushing the spleens through a 50-micron filter using magnesium- and calcium-free PBS. Cells were isolated using Dynabeads™ FlowComp™ Mouse CD8 Kit (Invitrogen, CAT#: 11462D) and treated with enough antibody for the number of cells isolated. Once T-cells were isolated, they were stained with eBioscience™ Cell Proliferation Dye eFluor™ 670 (100,000 cells per 500µL of 10µM solution) and plated in 96-well plates (100k cells/well) coated with 2µg/mL CD3e and 5µg/mL CD28 antibodies (LifeTech 16-0031-82 and 16-0281-82, respectively). DME-F12 SILAC media was prepared by adding leucine, lysine, arginine, pen/strep, and FBS, and supplemented with insulin/transferrin/selenium supplement and 100 ng/mL murine IL-2 (Sino Biologics). Enrichment of T cells was verified by flow cytometry with CD45- APC (BioLegend 103112), CD3e-PE (BioLegend 100308), CD4-PECy7 (BioLegend 100528) and CD8-FITC (BioLegend 100706). T-cells were plated in 96 well plates in media with methionine concentrations corresponding to the mouse they were isolated from (i.e. control group in regular methionine; MR group in low methionine). Neutrophils were also added at 100k cells/well for experiments requiring neutrophils. Cells were allowed to proliferate for 3 days before collecting and staining with CD3e-PE (BioLegend 100308) and CD11b- PECy7 (BioLegend 101216) antibodies. Proliferation dye staining intensity was measured in CD3+ cells using flow cytometry and analyzed via FlowJo.

### Neutrophil/Bone Marrow Isolation

Mice were sacrificed and the hind legs were removed to collect bone marrow (BM). Tissue was removed from bones of the leg that were placed in PBS without magnesium and calcium (used throughout) on ice. Bones were flushed with 1 mL of the PBS into a 1.5 mL tube. A histopaque gradient was created by adding 1.5 mL of histopaque 11191 into a FACs tube. The histopaque 10771 was slowly layered on top of histopaque 11191. Tilting the FACs tube can help to create this gradient slowly. Total BM was added to the top of the layers, very slowly. Tubes were spun in a centrifuge at 1000xg for 25 minutes at RT with no brake or accelerator. The top layer of the gradient was collected and placed in a 15 mL tube. The layer at the interface of the two gradients was also collected into the 15 mL tube. Cells were washed with 5 mL of the PBS and spun down at 1000xg for 5 minutes. Cells were resuspended in 1 mL of PBS and transferred to a 1.5 mL tube. All cells were counted.

### HALO® AI

All HALO® data analysis for immune cell subtypes utilized the macro version 33 that was previously described [50]. HALO® AI analysis concerning the tumor versus stroma was completed by first training the software to identify the difference between tumor and stroma. Once the program was efficient at differentiating the two, it was implemented to analyze all samples. The HALO® program was also used to set thresholds for IHC staining intensities, which was then analyzed for all slides by the program.

### Statistical Analysis

Statistical analyses were carried out using GraphPad Prism or Microsoft Excel. Unless otherwise stated, all numerical data are presented as mean ± standard error of the mean. Viral infections and selections were performed at least twice to produce distinct cultures to compare. For RNA-seq, one representative RNA sample was sent for each culture. For western blotting, samples from at least two distinct cell lines (different infection/selections or different passages of parental) were used to produce at least three western blots for each experiment. For statistical analysis, IC_50_ values were compared in GraphPad Prism by the sum-of-squares F statistic. For continuous data with sufficient n (>4) the parametric two tailed t-test or non-parametric Mann Whitney U tests were used depending on the result of a Shapiro-Wilks normality test. For n<4, a t-test was used. For two datasets, an outlier was removed according to ROUT method Q=1% in GraphPad Prism. For grouped analyses, one-way or two-way ANOVA were used and pairwise comparisons were made between specific groups. Holm-Šídák’s corrected p values or uncorrected Fisher’s LSD p values are reported as stated in figure legends. A p value of less than 0.05 was considered significant and p values just above that threshold were reported as exact numbers.

### Data Availability

RNA-sequencing data from this study are available at NCBI GEO database under accession number GSE270333. All other data are available upon reasonable request. TCGA data was obtained from NCI Genomic Data Commons to generate heat maps and gene expressions in transcripts per million (TPM) using the Legacy GRC37 dataset.

## ACKNOWLEDGEMENTS

The authors thank Dr. Dana Napier for assistance with the HALO® AI software. This work was supported in part by NCI R01 CA237643, NIGMS P20 GM121327-03, American Cancer Society Grant 133123-RSG-19-081-01-TBG, the American Institute for Cancer Research Grant 710410, the American Lung Association Innovation Award IA-046815, and the Markey Research Women Strong Distinguished Researcher Award 3048116064 (CFB), NCI T32 CA165990 (DRP), Markey STRONG Scholars Program through the American Cancer Society IRG-19-140-31 (EMS), and NIEHS T32 5T32ES007266 (TJD). This research was also supported by the Biostatistics & Bioinformatics, Cancer Research Informatics, Biospecimen Procurement & Translational Pathology, Flow Cytometry & Immune Monitoring Shared Resource Facilities, and pilot grant funding from the University of Kentucky Markey Cancer Center (P30 CA177558).

## AUTHOR CONTRIBUTIONS

KJN: Conceptualization, data curation, formal analysis, writing-original draft, writing-review and editing. XS: Investigation, data curation. ARC: Investigation, data curation, formal analysis. EMS: Investigation, data curation, formal analysis. ALB: Data curation. DPE: Investigation. CMG: Investigation, data curation, formal analysis. TJD: Investigation. DRP: Investigation writing-review and editing. AL: Investigation, data curation. JL: Conceptualization, data curation, formal analysis. CFB: Conceptualization, data curation, funding acquisition, investigation, writing-original draft, writing-review and editing.

