## Supplemental Tables and Figures for "Methionine Restriction Reduces Lung Cancer Progression and Increases Chemotherapy Response"

Kassanda J. Naughton et al.

### Supplementary Figures 1-7

### Supplementary Table 1

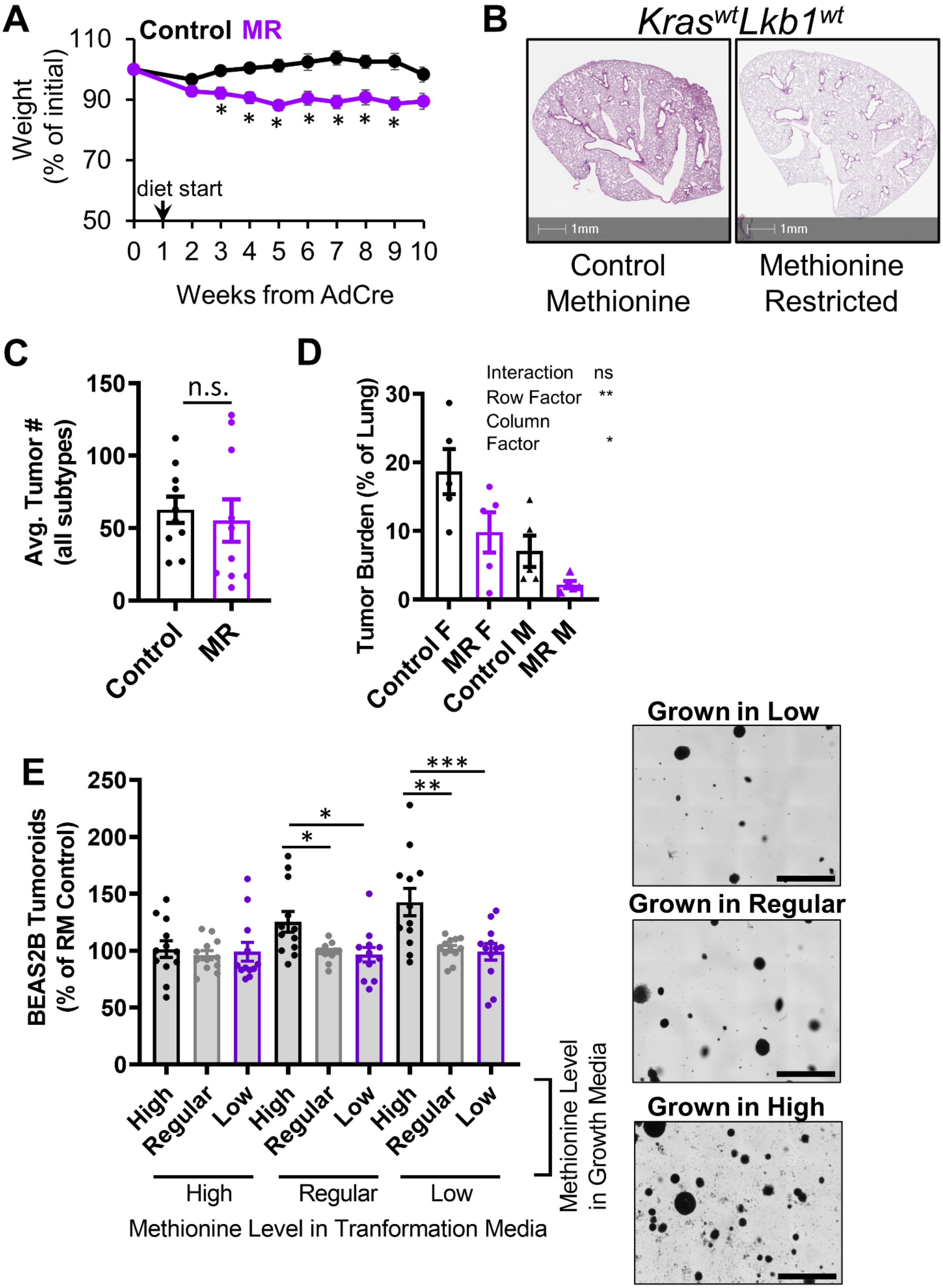


**Supplementary Figure 1:**

**A)** Average weight of mice on control or methionine restriction (MR) chow relative to starting weight, mean +/- SEM is plotted, n=10 mice per group, * indicates p<0.006 by two-tailed t-test. **B)** Representative H+E images of lungs of *Kras*-WT mice on control and methionine restricted diets, scale bars=1mm. **C)** Average tumor number of control and methionine restricted mice across all tumor subtypes plotted as mean +/- SEM, n=10 for each group, n.s.=not significant by two-tailed Mann-Whitney U test. **D)** Tumor burden of control and methionine restricted mice separated into groups based on sex plotted as mean +/- SEM, calculated based on the percent of total lung area that was identified as tumor. All groups n=5, interaction factor n.s.=not significant, Row Factor ** indicates p=0.0014 for male vs. female, Column Factor * indicates p=0.0144 for control methionine vs. methionine restriction by two-way ANOVA. **E)** BEAS-2B tumoroid counts after *KrasG12V* viral transformation in methionine media of different concentrations (High: 574.9 µM, Regular: 115.3 µM, Low: 57.9 µM) and grown in soft agar of different methionine media concentrations plotted as mean +/- SEM, n=12 for all groups, * indicates p<0.0434, ** p=0.0012, *** p=0.0006 by one-way ANOVA with Holm-Šídák's multiple comparisons test. Representative brightfield images of soft agar colonies transformed in regular methionine media and grown in low, regular, and high methionine media. Scale bars=2000µm.

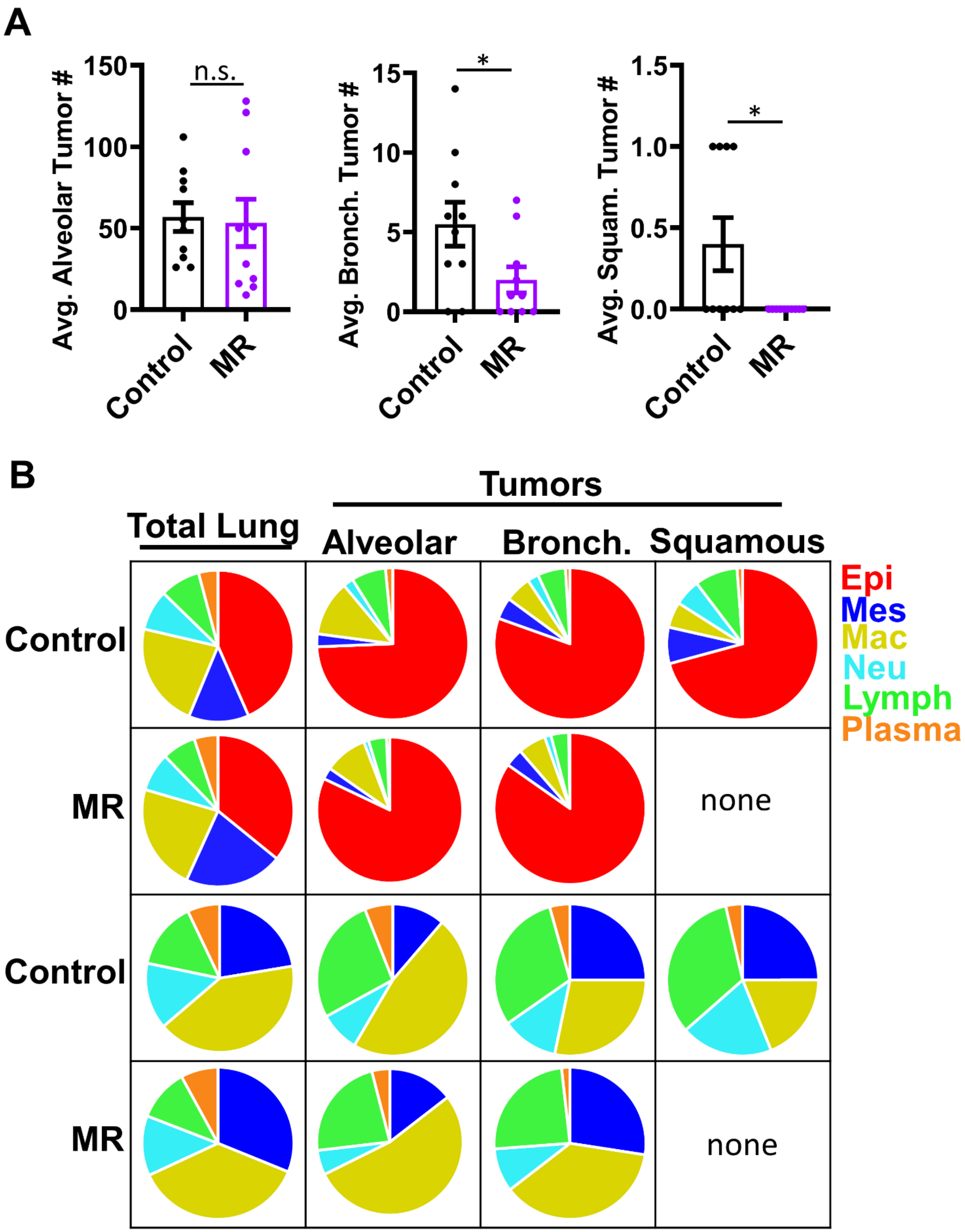


**Supplementary Figure 2:**

**A)** Average number of alveolar tumors in each control and methionine restricted mouse plotted as mean +/- SEM, n=10 mice per group, n.s.=not significant by two-tailed t-test. Average number of bronchiolar tumors in each control and methionine restricted mouse plotted as mean +/- SEM, n=10 mice for each group, * indicates p=0.0430 by two-tailed t-test. Average number of squamous tumors in each control and methionine restricted mouse plotted as mean +/- SEM, n=10 mice per group, * indicates p=0.0248 by two-tailed t-test. **B)** Pie charts of immune cells in control and methionine restricted mice separated by total lung, alveolar, bronchiolar, and squamous tumors. First rows of control methionine and MR include epithelial cells and second control methionine and MR exclude epithelial cells from percentages and instead compare all non-epithelial cells.

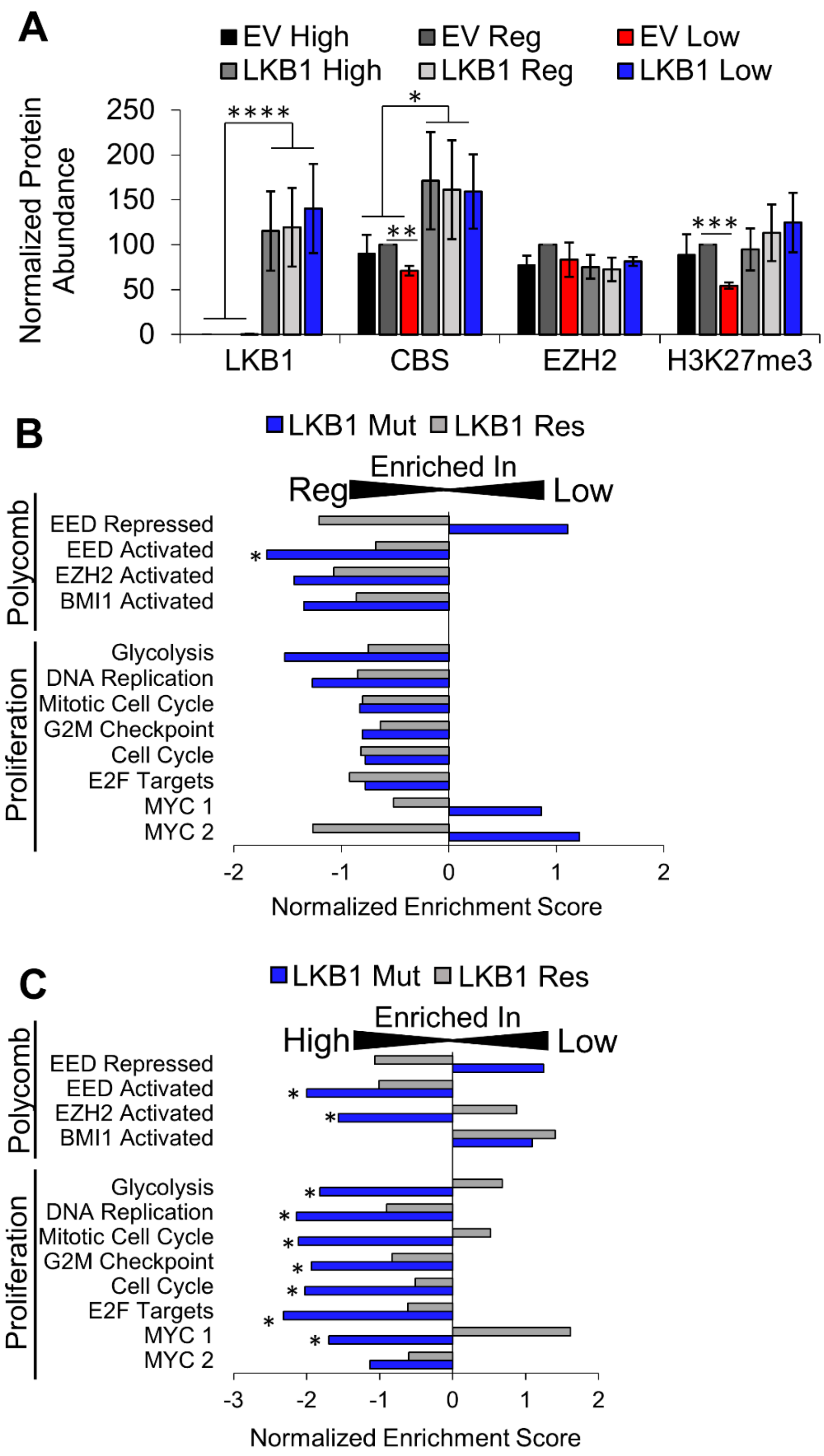


**Supplementary Figure 3:**

**A)** Western blots were quantified using the ImageJ Gel Analyzer tool on 16-bit, inverted grayscale images for A549 cells with and without LKB1 rescue in high, regular, and low methionine media. Values plotted as mean band intensity +/-SEM from 3 separate blots run from separate protein lysates. * indicates p=0.0104, **p=0.0059, ***p=0.0002, and ****p<0.0001 by two-tailed t-test. **B)** Gene Set Enrichment Analysis of pathways involved in Polycomb regulation and proliferation using the RNA sequencing data of A549 cells with and without LKB1 re-expression in regular versus low methionine RPMI media. * indicates $p<0.05$ using the FDR q-value. **C)** Gene Set Enrichment Analysis of pathways involved in Polycomb regulation and proliferation using the RNA sequencing data of A549 cells with and without LKB1 re-expression in high versus low methionine RPMI media. * indicates $p<0.05$ using the FDR q-value.

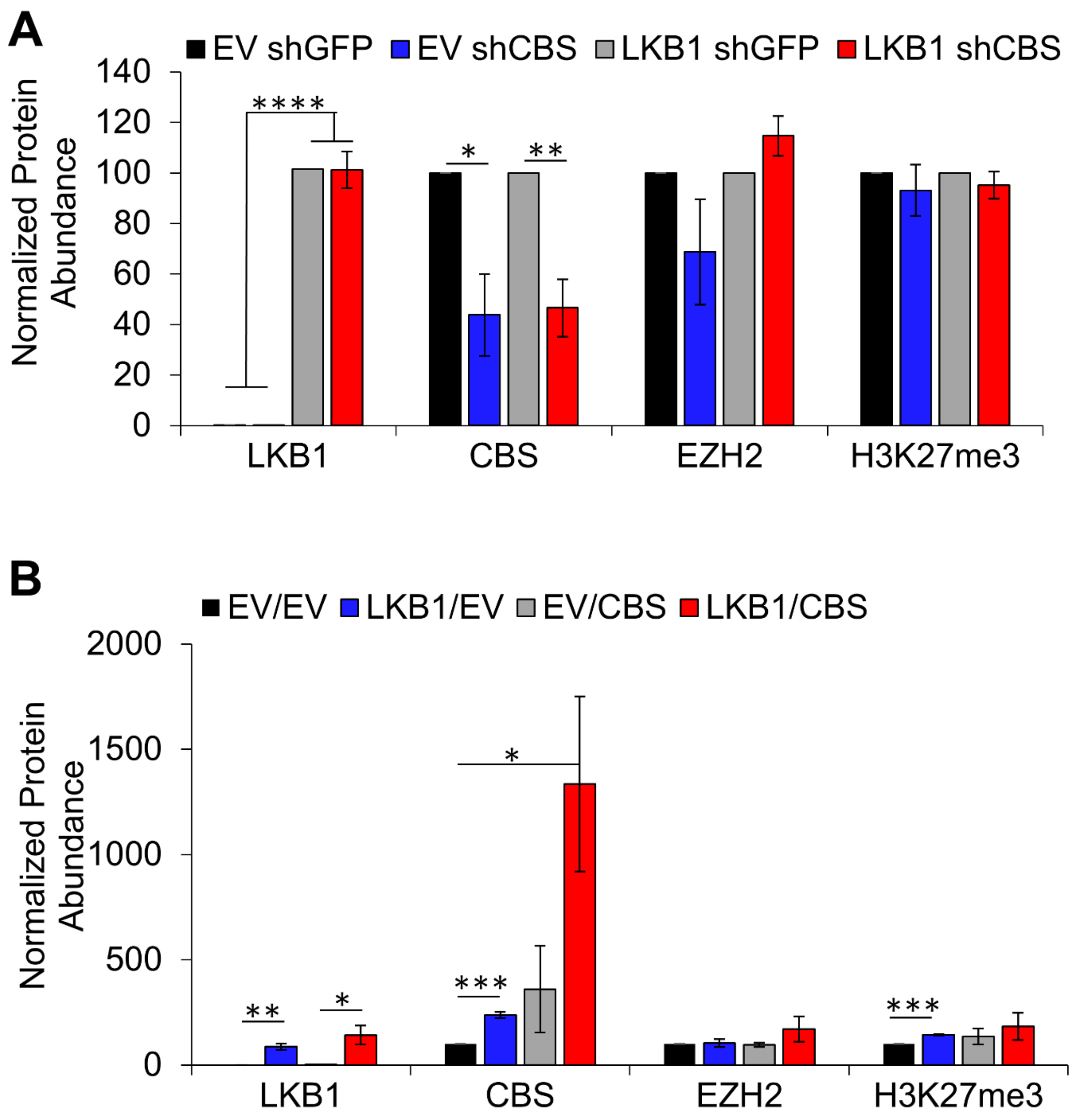


**Supplementary Figure 4:**

**A)** Western blots were quantified using the ImageJ Gel Analyzer tool on 16-bit, inverted grayscale images for A549 cells with and without LKB1 rescue and with and without CBS knockdown. Values plotted as mean band intensity +/-SEM from 4 separate blots run from separate protein lysates. * indicates p=0.0134, **p=0.0032, and ****p<0.0001 by two-tailed t-test. **B)** Western blots were quantified using the ImageJ Gel Analyzer tool on 16-bit, inverted grayscale images for A549 cells with and without LKB1 rescue and with and without CBS overexpression. Values plotted as mean band intensity +/-SEM from 3 separate blots run from separate protein lysates. * indicates p<0.0411, **p=0.0043, and ***p=0.0006 by two-tailed t-test.

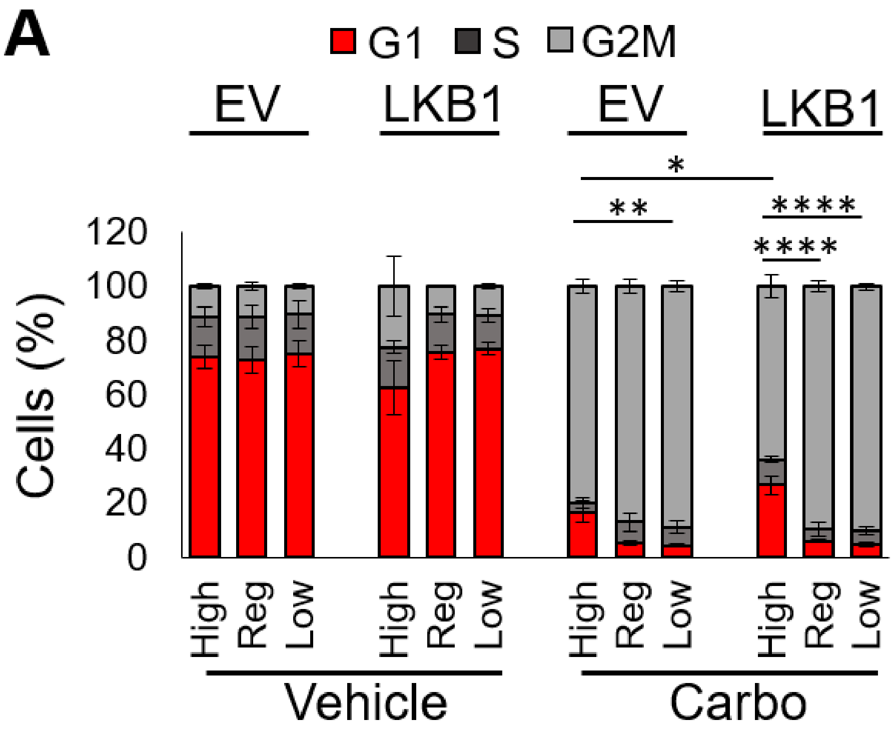


**Supplementary Figure 5:**

**A)** 7AAD cell cycle flow cytometry on A549 cells with and without LKB1 re-expression grown in high, regular, and low methionine media with and without carboplatin treatment at 60 µM, plotted as mean +/- SEM, n=4 biological replicates. Statistics are shown for G1 phase comparing carbo groups by one-way ANOVA. * indicates p=0.0329, **p=0.0072, and ****p<0.0001 by two-tailed t-test.

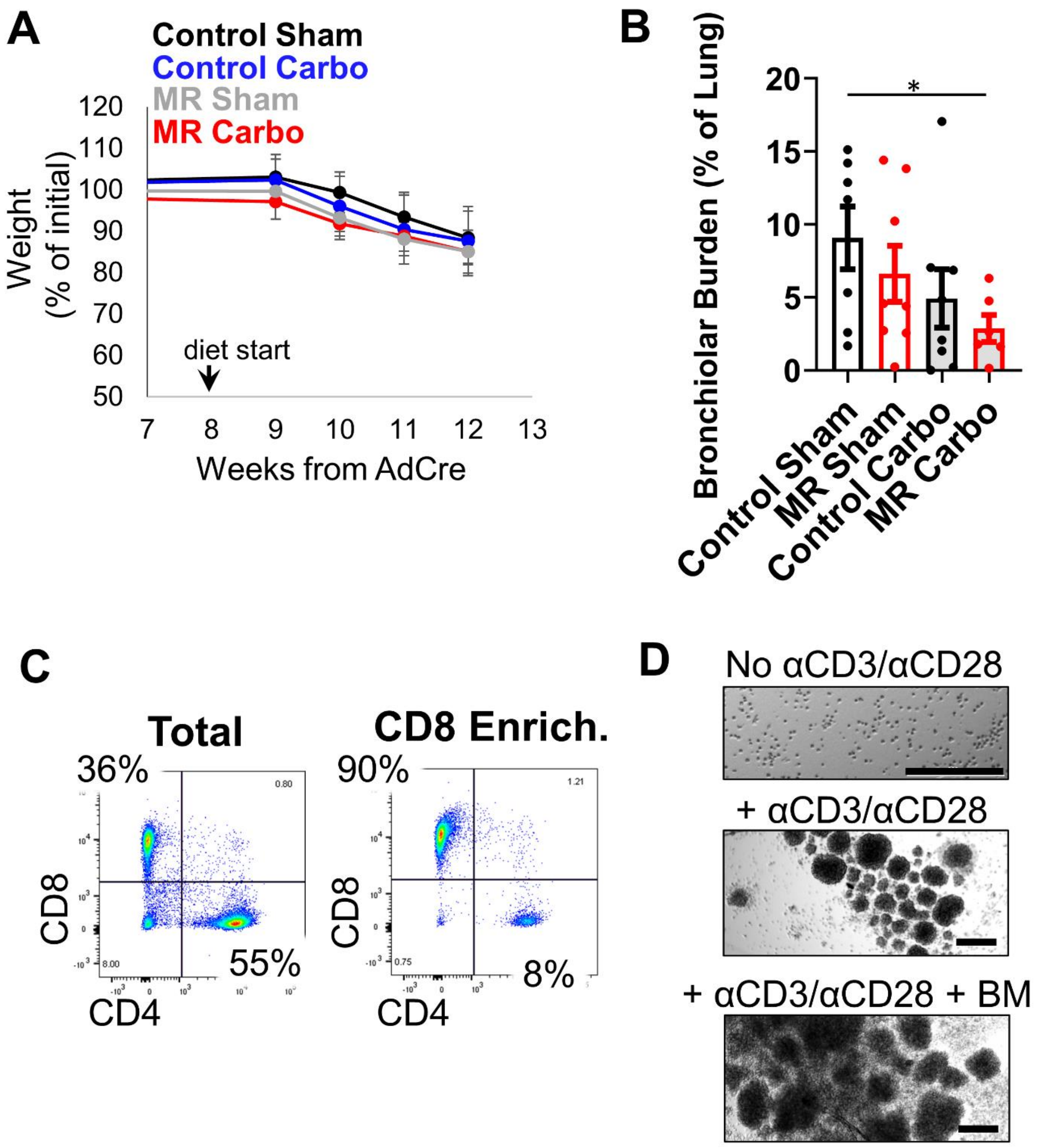


**Supplementary Figure 6:**

**A)** Weights of mice on the carboplatin and methionine treatment study from diet start to end of study. Plotted as mean +/- SEM as percentage of initial weight. **B)** Bronchiolar tumor burden calculated as percent of total lung that was identified as bronchiolar tumor, plotted as mean +/-SEM, n=7 for control sham, n=8 for MR sham, n=8 for control carbo, and n=6 for MR carbo. *indicates p=0.0384 by one-way ANOVA. **C)** T Cell isolation validation by flow cytometry, gated on CD45+/CD3+ from total spleen before and after CD8+ isolation. **D)** Images of T cells in culture with and without activation and/or co-culturing with cells collected from total bone marrow, scale bar=200µm.

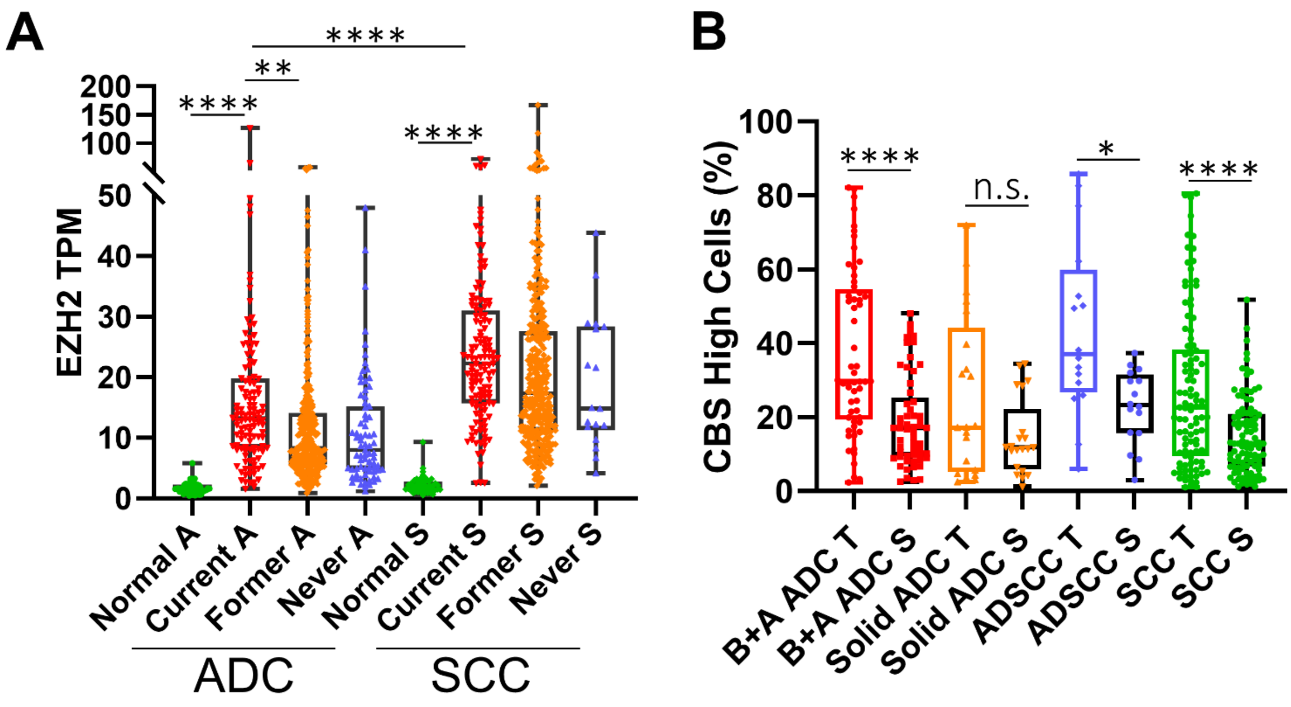


**Supplementary Figure 7:**

**A)** Human TCGA data from patient samples grouped by smoking status (normal tissue, current, former, and never smokers) and by tumor type (ADC=adenocarcinoma, SCC=squamous cell carcinoma) for *EZH2* expression, plotted as mean+/-SEM in transcripts per million (TPM), n=58 for normal ADC, n=120 for current ADC, n=307 for former ADC, n=75 for never ADC, n=51 for normal SCC, n=133 for current SCC, n=338 for former SCC, and n=18 for never SCC. **indicates p=0.0082, ****p<0.0001 by one-way ANOVA using Holm-Šídák's multiple comparisons test. **B)** Human tumor microarray data from patient samples grouped by tumor type (B+A ADC = bronchiolar + alveolar adenocarcinoma, ADSCC = adenosquamous cell carcinoma) and by tumor (T) and stroma (S) measurements for high *CBS* expression, plotted as mean+/-SEM in transcripts per million (TPM), n=57 for B+A ADC T and B+A ADC S, n=21 for Solid ADC T and Solid ADC S, n=16 for ADSCC T and ADSCC S, and n=102 for SCC T and SCC S. *indicates p=0.0124, ****p<0.0001 by one-way ANOVA using Holm-Šídák's multiple comparisons test.

**Supplementary Table 1: Genotyping Primer**

| **Lkb1 floxed Primers** | **Sequence** |
|---|---|
| Primer 1 | 5’- GGG CTT CCA CCT GGT GCC AGC CTG T -3’ |
| Primer 2 | 5’- GAG ATG GGT ACC AGG AGT TGG GGC T -3’ |
| Primer 3 | 5’- TCT AAC AAT GCG CTC ATC GTC ATC CTC GGC -3’ |
| Expected Bands | WT = 220bp, floxed=300bp |
| **Kras LSL Primer** | **Sequence** |
| Primer 1 | 5’ - GTC TTT CCC CAG CAC AGT GC - 3’ |
| Primer 2 | 5’ - CTC TTG CCT ACG CCA CCA GCT C– 3’ |
| Primer 3 | 5’ – AGC TAG CCA CCA TGG CTT GAG TAA GTC TGC A– 3’ |
| Expected Bands | WT=622bp, LSL intact=500bp, LSL-excised=650bp |
